# ZNF397 Loss Triggers TET2-driven Epigenetic Rewiring, Lineage Plasticity, and AR-targeted Therapy Resistance in AR-dependent Cancers

**DOI:** 10.1101/2023.10.24.563645

**Authors:** Yaru Xu, Zhaoning Wang, Martin Sjöström, Su Deng, Choushi Wang, Nickolas A Johnson, Julisa Gonzalez, Xiaoling Li, Lauren A Metang, Carla Rodriguez Tirado, Atreyi Mukherji, Garrett Wainwright, Xinzhe Yu, Yuqiu Yang, Spencer Barnes, Mia Hofstad, Hong Zhu, Ariella Hanker, Housheng Hansen He, Yu Chen, Zhao Wang, Ganesh Raj, Carlos Arteaga, Felix Feng, Yunguan Wang, Tao Wang, Ping Mu

## Abstract

Cancer cells exhibit phenotypical plasticity and epigenetic reprogramming, which allows them to evade lineage-dependent targeted treatments by adopting lineage plasticity. The underlying mechanisms by which cancer cells exploit the epigenetic regulatory machinery to acquire lineage plasticity and therapy resistance remain poorly understood. We identified Zinc Finger Protein 397 (ZNF397) as a *bona fide* co-activator of the androgen receptor (AR), essential for the transcriptional program governing AR-driven luminal lineage. ZNF397 deficiency facilitates the transition of cancer cell from an AR-driven luminal lineage to a Ten-Eleven Translocation 2 (TET2)-driven lineage plastic state, ultimately promoting resistance to therapies inhibiting AR signaling. Intriguingly, our findings indicate that TET2 inhibitor can eliminate the AR targeted therapies resistance in ZNF397-deficient tumors. These insights uncover a novel mechanism through which prostate and breast cancers acquire lineage plasticity via epigenetic rewiring and offer promising implications for clinical interventions designed to overcome therapy resistance dictated by lineage plasticity.

**Statement of Significance:** This study reveals a novel epigenetic mechanism regulating tumor lineage plasticity and therapy response, enhances understanding of drug resistance and unveils a new therapeutic strategy for prostate cancer and other malignancies. Our findings also illuminate TET2’s oncogenic role and mechanistically connect TET2-driven epigenetic rewiring to lineage plasticity and therapy resistance.

## Introduction

Emerging evidence has identified cellular plasticity, specifically lineage plasticity, as a major hallmark of cancer ^1^. Cancer cells possess the ability to alter their established lineage by reverting to a stem-like state and subsequently redifferentiating into an alternative lineage, thereby evade therapies targeting their original lineage-directed survival programs ^2–5^. This plasticity has been demonstrated to cause resistance to many standard cancer therapies in various types of human cancers, including prostate, breast, lung, and pancreatic cancers, as well as melanoma ^3,5–11^. Although numerous genomic aberrations have been associated with the acquisition of lineage plasticity ^3,5,6,8,12–19^, only a limited number of tumors exhibit these known alterations ^20^, suggesting that a significant proportion of patients experience resistance due to lineage plasticity through unidentified mechanisms. More importantly, the underlying mechanism by which cancer cells initiate the lineage transition to a plastic state remains largely unclear. Lastly, no effective therapies currently exist to target lineage plasticity-driven resistance in tumors, emphasizing the urgent need to identify key modifiers of lineage plasticity and, consequently, druggable targets to reverse it.

In addition to genomic changes, non-mutational epigenetic reprogramming has been identified as a crucial factor in cellular lineage decisions and therapy response in various cancers^21^. This includes epigenetic modifications mediated by BET family proteins, EZH2, CHD1, LSD1, and the SWI/SNF complex ^6,8,22–27^. Among the many important epigenetic modifiers associated with lineage regulation, one member of the Ten-Eleven Translocation (TET) proteins, the methylcytosine dioxygenase TET2, has a key role in regulating cell fate and lineage decisions during embryonic development and carcinogenesis ^28–33^. Importantly, both TET2 expression and TET2-dependent oxidation of methylated cytosine (5mC) to 5-hydroxymethylcytosine (5hmC) are repressed by Androgen Receptor (AR) signaling, which is the luminal epithelial lineage-specific survival factor and drives the growth of primary prostate cancer (PCa) ^34–36^. In contrast, TET2 expression and 5hmC levels are functionally enriched during the development of stem-like and neuronal lineages ^33,37–44^, which are two alternative lineages known to confer AR-targeted therapy resistance in PCa. However, the exact molecular function of TET2 in mediating lineage plasticity, as well as the mechanism by which cancer cells hijack the TET2-mediated epigenetic regulation machinery to acquire lineage plasticity and therapy resistance, remains poorly understood.

Here, we have identified Zinc Finger Protein 397 (ZNF397) as a key co-activator of the AR, which is necessary to maintain both AR signaling and the luminal lineage in prostate and breast cancers. In contrast, ZNF397 interacts with and blocks TET2 binding to many of the TET2 target genes. The frequent loss of ZNF397 in prostate cancer is a driving event that promotes the transition of cancer cells from an AR-driven luminal lineage to a TET2-driven lineage plastic state expressing epithelial-to-mesenchymal transition (EMT)-like, stem-like and neuroendocrine (NE)-like lineages, rendering cells unresponsive to AR-targeted therapies. Contrary to the conventional view that TET2 functions as a tumor suppressor in primary PCa, we demonstrated that TET2 is a crucial mediator of lineage plasticity-driven therapy resistance in metastatic castration resistant prostate cancer (mCRPC). Both genetic and pharmacological inactivation of TET2 impeded the resistance to AR targeted therapies in ZNF397-deficient tumors. Collectively, these findings uncover a cell-intrinsic molecular switch that dictates tumor lineage plasticity and suggest that targeting TET2 has the potential to overcome AR therapy resistance.

## Results

### ZNF397-deficiency promotes PCa tumorigenesis and resistance to AR-targeted therapy

Zinc finger proteins constitute the most extensive family of transcription factors, functioning as either transcriptional repressors or activators for numerous downstream genes that play critical roles in diverse physiological and carcinogenic processes. ZNF397, a member of the classical Cys_2_His_2_ group of SCAN-zinc-finger proteins, was initially identified as a mammalian centromere protein essential for centromere localization ^45^. Although ZNF397 depletion (both deep and shallow) has been frequently observed in various cancers (up to 60% of patients, Fig.S1A), especially hormone-driven cancers such as prostate and breast cancers, its role in tumorigenesis remains elusive. Interestingly, loss of ZNF397 emerged as one of the top candidate events responsible for AR-targeted therapy resistance (Fig.1A), based on a re-analysis of our previous *in vivo* library screening using the advanced ranking algorithm MAGeCK ^8^. Notably, a Pearson correlation analysis of a genomic landscape study of metastatic castration-resistant prostate cancer (mCRPC, SU2C) showed that *ZNF397* expression level is significantly correlated with progression-free survival time of PCa patients on AR-targeted therapies (Fig.1B) ^8,20^. To further dissect this correlation, we divided the cohort into two groups based on ZNF397 expression (above or below median expression) and observed that the patients with low ZNF397 expression developed AR-targeted therapy resistance significantly faster than the patients with high ZNF397 expression (Fig.1C). Furthermore, the patients with low ZNF397 expression had a significantly increased “hazard to develop therapy resistance” metric compared to patients with high ZNF397 expression, based on a Cox proportional hazards ratio analysis (Fig.1D). The correlation between low ZNF397 expression and poor clinical outcome, including shorter progression-free survival and overall survival, were also observed in breast, ovarian and lung cancers (Fig.S1B-G) ^46–48^. Collectively, the clinical correlations between ZNF397 expression and resistance to targeted therapies raise the intriguing possibility that this understudied zinc finger protein, ZNF397, plays a key role in mediating tumorigenesis and therapy resistance of PCa and other cancers.

**Figure 1:**
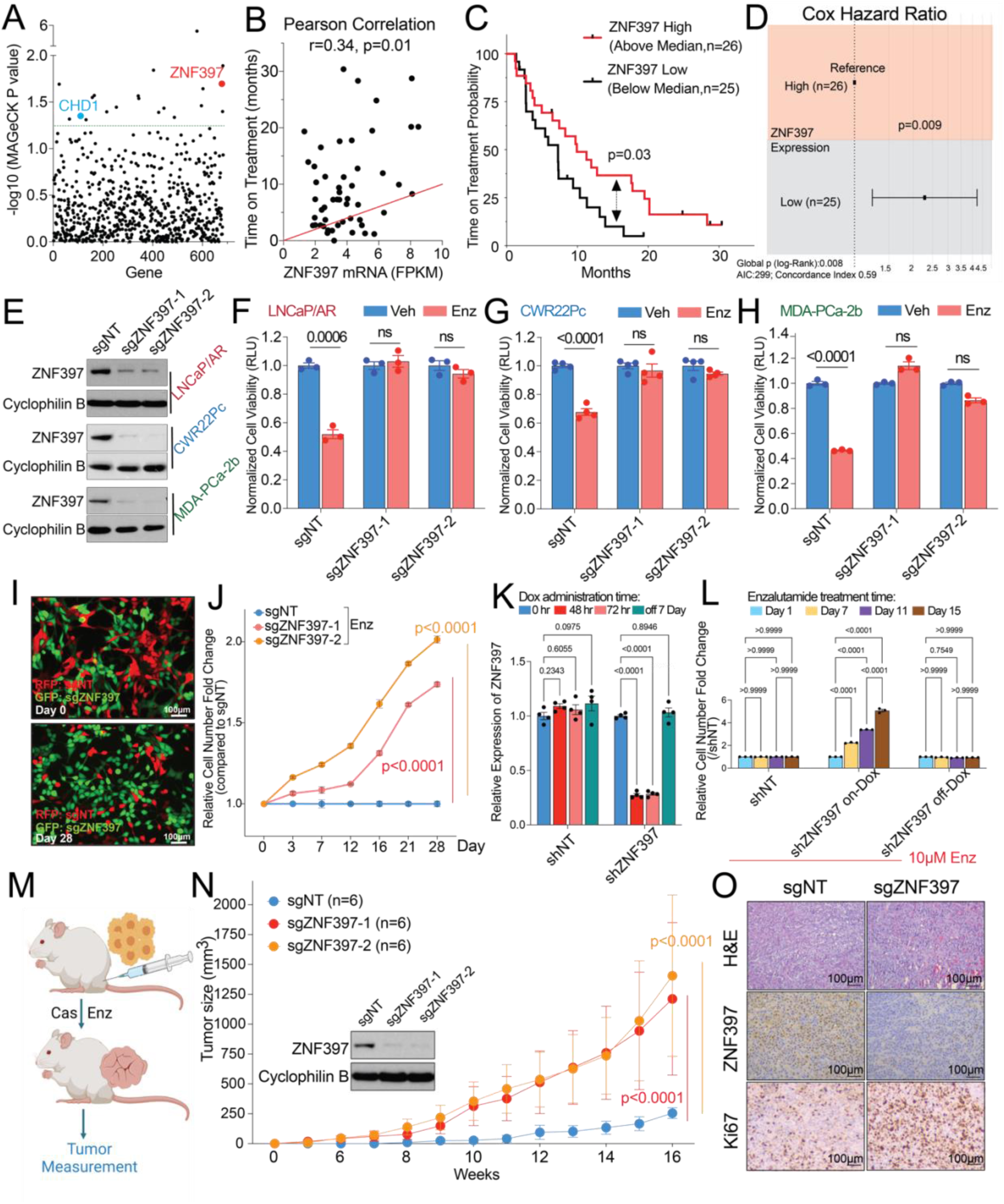
ZNF397 deficiency promotes PCa tumorigenesis and resistance to AR-targeted therapy. (A) Dot plot represents the results of MAGeCK algorithm-based re-analysis of the *in vivo* library screening. ZNF397 is presented as red dot and CHD1 (previously identified top hit) is presented as blue dot. P values were calculated with MAGeCK and green line represents the statistical significance cutoff line (p<0.05). (B) Pearson correlation analysis shows the relationship between ZNF397 mRNA and time of treatment on Abi/Enz/Apa in the SU2C mCRPC patient cohort. Abi: abiraterone, Enz: enzalutamide, Apa: apalutamide. (C) Kaplan-Meier curve represents the treatment duration on AR-targeted therapies of patients with high (above median, n=26) or low (below median, n=25) expression of ZNF397 in a prostate cancer patient cohort. The p value was calculated with the log-rank test. (D) Cox proportional hazard ratio analysis of patients with high (above median, n=26) or low (below median, n=25) expression of ZNF397 in this cohort. The p value was calculated with the log-rank test. (E) Western blot of ZNF397 in a series of human PCa cell lines with CRISPR-based ZNF397-KO. (F-H) Bar plots represent the relative cell viability of LNCaP/AR (F), CWR22Pc (G), and MDA-PCa-2b cells (H) transduced with Cas9 and annotated CRISPR guide RNAs, measured as values of relative luminescence units (RLU) and normalized to vehicle-treated conditions. Enz denotes enzalutamide treatment (10 μM for LNCaP/AR and MDA-PCa-2b, 1 μM for CWR22Pc) for several days (7 days for LNCaP/AR, 5 days for CWR22Pc and MDA-PCa-2b). Veh denotes DMSO. (I) Fluorescence microscope imaging shows the cell mixtures of sgZNF397 cells (green) and sgNT cells (red) on day 0 and day 28 of the FACS-based competition assay cultured with 10 μM Enz in CSS medium; representative pictures of n = 3 independently treated cell cultures were shown. (J) Relative cell number fold change of LNCaP/AR cells transduced with Cas9 and annotated CRISPR guide RNAs, measured by FACS-based competition assay. Enz denotes 10 µM enzalutamide treatment for 28 days. (K) Relative gene expression levels of ZNF397 in the inducible shZNF397 LNCaP/AR cells treated with Dox for various lengths of time. Data are normalized to 0 h. (L) Relative cell number fold change of LNCaP/AR cells transduced with inducible shZNF397. On-Dox cells were consistently exposed to Dox during the enzalutamide treatment period, while Off-Dox cells were treated with Dox for 3 days before removal for 7 days. Data are normalized to Day 1, measured by FACS-based competition assay. Enz denotes 10 µM enzalutamide treatment for 15 days. (M-N) Tumor growth curve of xenografted LNCaP/AR cells transduced with annotated guide RNAs in castrated mice. Enz denotes enzalutamide treatment at 10 mg/kg from day 1 of grafting. Cas denotes castration. The number of tumors in each group was annotated. (O) H&E and immunohistochemical staining of ZNF397 and Ki67 on sgNT and sgZNF397 xenograft tumor slides. Scale bar represents 100 μm. For all panels unless otherwise noted, 3 biological replicates in each group and mean ± s.e.m. is represented. p values were calculated using two-way ANOVA with Bonferroni multiple-comparison test. See also **Figure. S1-2**.

To investigate the potential role of ZNF397 in mCRPC, we first knocked-out (KO) ZNF397 from a panel of PCa cell line models that are sensitive to the AR antagonist enzalutamide (Enz), including LNCaP/AR, CWR22Pc, and MDA-PCa-2b cells, using multiple independent stable CRISPR/Cas9 guide RNAs (Fig.1E). Consistent with clinical observations, ZNF397-KO led to significant enzalutamide resistance in all three cell line models, as measured by cell viability assay (Fig.1F-H) and FACS-based competition assays (Fig.1I, J, Fig.S2A, B). To further dissect the dynamics of this resistance, we utilized a doxycycline (Dox)-inducible shRNA system to knockdown (KD) ZNF397 and observed that the ZNF397-KD conferred enzalutamide resistance was both rapid and reversible (Fig.1K, L), suggesting that ZNF397 may mediate enzalutamide response through transcriptional or epigenetic mechanisms. Furthermore, ZNF397 deletion also led to significant resistance to AR therapies *in vivo*, as shown by the growth of LNCaP/AR xenografted tumors in castrated mice (Fig.1M, N). Immunohistochemistry staining confirmed that ZNF397-KO tumors had substantially increased Ki67 signal compared to control tumors (Fig.1O), indicating that loss of ZNF397 protects prostate cancer tumors from therapy-induced inhibition of proliferation. Notably, ZNF397-KO tumors did not progress at a faster rate than control tumors in castrated mice treated with vehicle (Fig. S2C) or intact mice treated with vehicle (Fig.S2D), suggesting a specific role of ZNF397 in mediating the response to AR-targeted therapies, rather than regulating general tumor cell proliferation.

### ZNF397 is required for the canonical AR-driven transcriptional program

To decipher the mechanism through which ZNF397-KO promotes resistance to the AR antagonist, enzalutamide, we first examined the expression of canonical AR target genes. Surprisingly, we observed persistent inhibition of AR signaling in enzalutamide-treated ZNF397-KO tumors (Fig.2A), contrary to the conventional resistance mechanism of AR signaling restoration in many resistant mCRPC tumors. Strikingly, the expression of canonical AR target genes was already downregulated in ZNF397-KO cells even prior to the treatment of enzalutamide (Fig.2B), which was validated by downregulation of the protein levels of AR target genes (Fig.2C). These results suggest that ZNF397-KO may lead to the loss of the AR transcriptional program and activate alternative transcriptional programs that relieve PCa from its dependency on AR signaling. Consistent with this hypothesis, AR agonist dihydrotestosterone (DHT)-induced upregulation of AR signaling was diminished in ZNF397-KO cells compared to control cells (Fig.2D), suggesting that ZNF397 is required for maintaining AR signaling. This hypothesis is further substantiated by the observation that ZNF397-KO protected the PCa cells from the cellular toxicity of excessive AR signaling activation by high dosages of DHT (Fig.2E).

**Figure 2:**
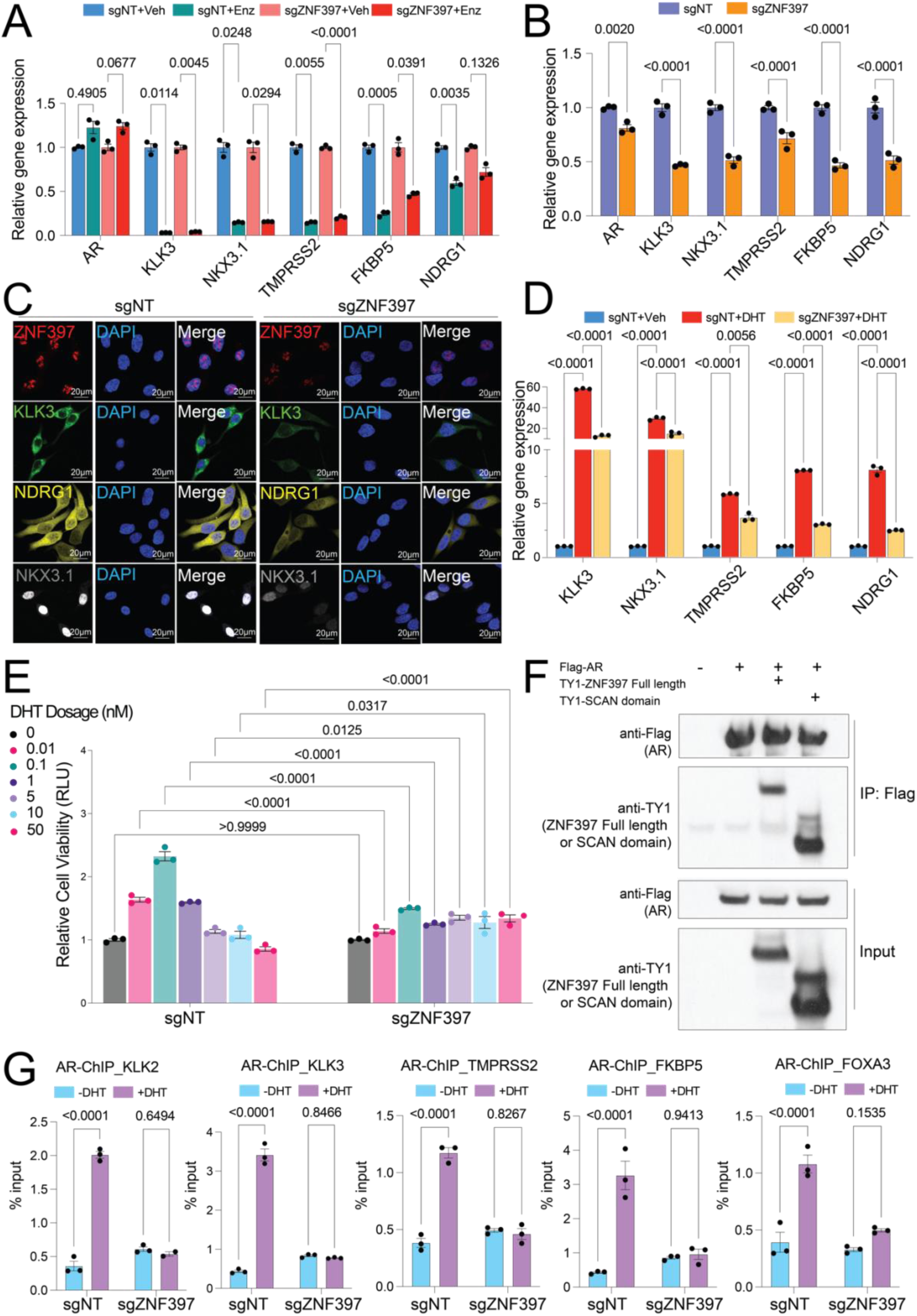
ZNF397 is required for the canonical AR-driven transcriptional program. (A) Relative gene expression levels of androgen receptor (AR) and AR target genes in LNCaP/AR cells transduced with Cas9 and annotated guide RNAs, and treated with vehicle (Veh, DMSO) or enzalutamide (Enz, 10 µM) for 7 days, normalized and compared to sgNT+Veh group. (B) Relative gene expression levels of AR and AR target genes in LNCaP/AR cells transduced with Cas9 and annotated guide RNAs, normalized and compared to sgNT+Veh group. p values were calculated using multiple t-tests with Benjamini correction. (C) Representative immunofluorescence staining images of LNCaP/AR cells transduced with Cas9 and annotated guide RNAs with annotated antibodies; n = 3 independent treated cell cultures are shown. (D) Relative gene expression levels of AR target genes in LNCaP/AR cells transduced with Cas9 and annotated guide RNAs, and treated with vehicle (Veh, DMSO) or 10 nM dihydrotestosterone (DHT) for 12 hrs, normalized and compared to sgNT+Veh group. (E) Bar plot representing the relative cell viability of LNCaP/AR cells transduced with annotated guide RNAs and treated with different doses of DHT for 7 days, measured as values of relative luminescence units (RLU) and normalized to vehicle-treated conditions. (F) Co-immunoprecipitation (Co-IP) of AR (with Flag tag) and full length or the SCAN domain of ZNF397 (with TY1 tag) in HEK293T cells. (G) AR ChIP-qPCR of the genomic loci of canonical AR target genes in LNCaP/AR cells transduced with annotated constructs, treated with Veh (-DHT) or DHT (+DHT). For all panels unless otherwise noted, n = 3 independently treated cell cultures and mean ± s.e.m. is represented. p values were calculated using two-way ANOVA with Bonferroni multiple-comparison test. See also **Figure.S3**

To further dissect the mechanism leading to impaired AR signaling, we performed coimmunoprecipitation (co-IP) assays and observed robust interactions of AR with both the full length and the SCAN domain of ZNF397 (Fig.2F), suggesting that ZNF397 maybe an essential cofactor for AR-driven transcription. Consistent with this hypothesis, AR Chromatin Immunoprecipitation (ChIP) analysis revealed that ZNF397-KO largely abolished AR binding to its canonical target genes (Fig.2G). Additionally, histone marker ChIP-qPCR analyses revealed that the knockout (KO) of ZNF397 considerably impairs the transcriptional activation of canonical AR target genes (Fig.S3). This finding is supported by the observed reduction in active histone markers, such as H3K27 acetylation (H3K27ac), H3K4 trimethylation (H3K4me3), and H3K4 dimethylation (H3K4me2) (Fig. S3A-C), as well as the increase in the repressive histone marker H3K27 trimethylation (H3K27me3) (Fig. S3D). Collectively, these results support the essential role of ZNF397 in maintaining the AR transcriptome in PCa cells.

### ZNF397-deficiency promotes lineage plasticity and multilineage transcriptional programs

Given that the shift to a lineage plastic and AR-independent state is a major mechanism underlying resistance to AR-targeted therapy, we conducted an analysis of the expression of canonical lineage marker genes in LNCaP/AR cells. We observed significant downregulation of AR-dependent luminal lineage genes and upregulation of lineage plastic genes, including basal, EMT-like, stem-like, and NE-like marker genes, in ZNF397-KO cells (Fig.3A). Immunofluorescence (IF) staining confirmed the downregulation of AR target gene (TMPRSS2) and luminal marker protein (KRT8), and upregulation of lineage plastic marker proteins (KRT14, VIM, CDH2, ASCL1) (Fig.3B, Fig.S4A). This transition from a luminal lineage solely driven by AR to an AR-independent multilineage state was further confirmed in another two human PCa models, namely, the CWR22Pc and MDA-PCa-2b cells (Fig.S4B, C).

**Figure 3:**
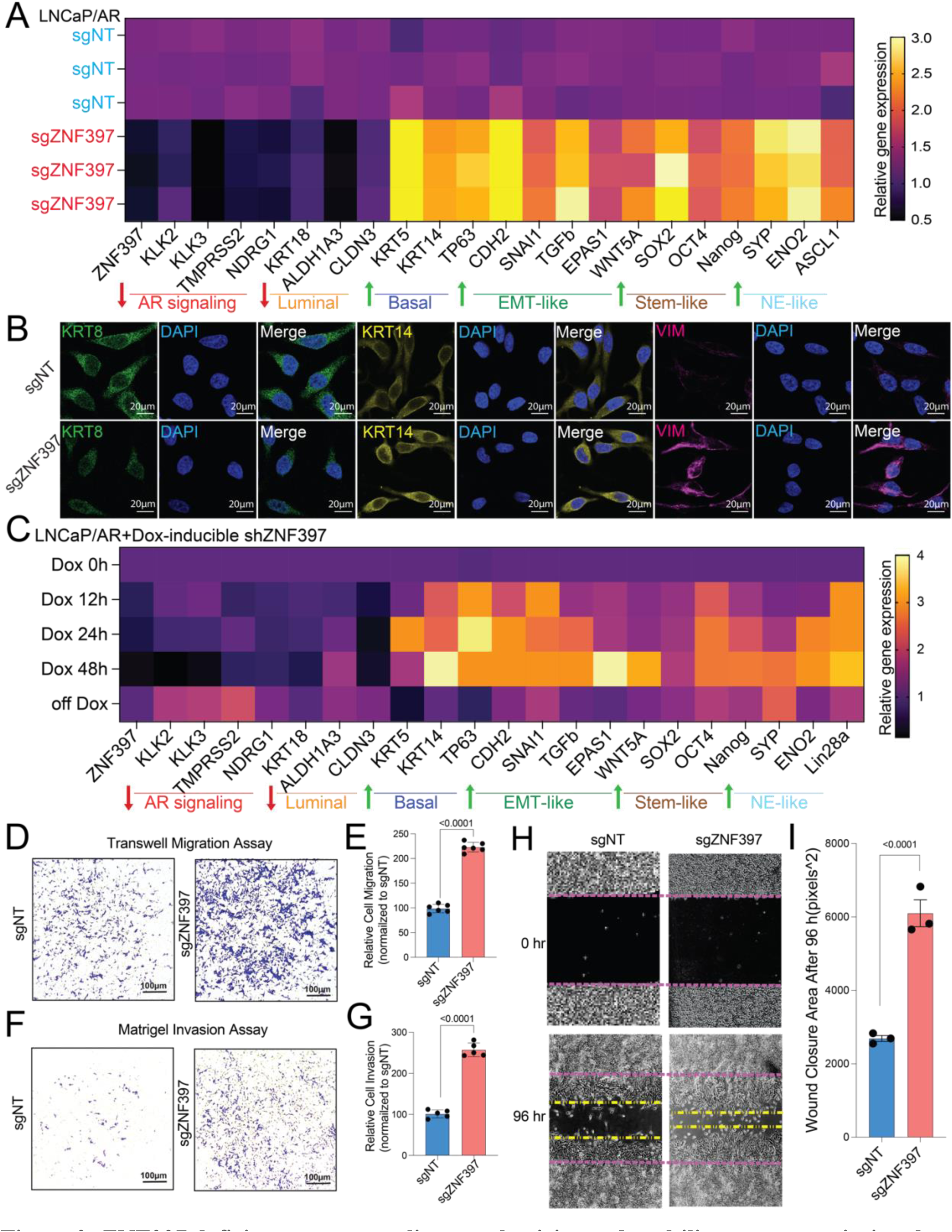
ZNF397 deficiency promotes lineage plasticity and multilineage transcriptional programs. (A) Relative gene expression levels of canonical lineage-specific marker genes in LNCaP/AR cells transduced with Cas9 and annotated guide RNAs, as measured by qPCR assay. Data are normalized to the average expression in sgNT cells. (B) Immunofluorescence staining of LNCaP/AR cells transduced with Cas9 and annotated guide RNAs using antibodies against lineage-specific markers; representative images from n = 3 independent treated cell cultures are shown. (C) Relative gene expression levels of canonical lineage-specific marker genes in inducible shZNF397 LNCaP/AR cells treated with Dox for varying lengths of time. Data are normalized to 0 h. (D) Representative images of an LNCaP/AR cell transwell migration assay from six independently treated cell cultures. (E) Quantification of migrated cell numbers from six representative images, taken from three independently treated cell cultures for each of the cell lines. (F) Representative images of an LNCaP/AR cell invasion assay from five independently treated cell cultures. (G) Quantification of the numbers of invading cells from five representative images, taken from three independently treated cell cultures for each of the cell lines. (H) Representative images of an LNCaP/AR cell wound healing assay from three independently treated cell cultures. (I) Quantification of the occupied areas of invading cells from three representative images, taken from three independently treated cell cultures for each of the cell lines. Unless otherwise noted, p-values were calculated using a two-tailed t-test. See also **Figures S1, S4** and **S5**.

To further decipher the dynamic of the lineage plasticity conferred by ZNF397-KO, we examined the expression of lineage marker genes in the doxycycline-inducible ZNF397-KD cells and revealed that the loss of luminal lineage program and acquisition of lineage plastic programs occurred within 12 hours upon Dox administration and KD of ZNF397 (Fig.3C). Remarkably, the expression of these lineage plastic programs was fully reversed to wildtype levels upon restoration of ZNF397 expression following Dox removal (Fig.3C), providing evidence for the reversibility of this phenotype. These results indicate that ZNF397 may mediate lineage plasticity and therapy resistance through transcriptional or epigenetic mechanisms, which is fast and reversible. Consistent with the induction of EMT-like lineage program, ZNF397-KO cells exhibited significantly increased migration and invasiveness compared to control cells, as demonstrated by various migration and invasion assays (Fig.3D-I). Finally, ZNF397-KO significantly promotes the prostasphere formation ability of tumor cells (Fig.S4D, E), which corroborates the acquisition of stem-like lineage program in ZNF397-KO cells.

Interestingly, the crucial role of ZNF397 in mediating AR transcription and lineage plasticity is not limited to prostate cancer. ZNF397-KO in three AR-positive breast cancer cell lines, MDA-MB-453, MFM-223 and SUM-185PE ^49^, led to enhanced tumor cell growth (Fig.S5A, B) and transition from an AR-driven luminal state to a lineage plastic and multilineage state (Fig.S5C-E). These results are consistent with the clinical correlation between low ZNF397 expression and shorter survival in breast cancer (Fig.S1B-C).

### Integrated analysis of ChIP-Seq, RNA-Seq and CRISPR screening reveals key drivers of lineage plastic programs

Given that ZNF397-KO eliminates AR binding and elevates the expression of genes associated with lineage plasticity, we hypothesized that the loss of ZNF397 activates alternative transcriptional programs that relieve prostate cancer tumors from their dependence on AR signaling and luminal lineage, thereby conferring resistance to AR-targeted therapy. To determine these alternative lineage-plastic transcriptional programs, AR ChIP-seq and RNA-seq were performed to assess the global changes in the cistromic and transcriptomic landscapes induced by ZNF397-KO, respectively. Globally, AR ChIP-Seq revealed a significant loss of AR binding peaks (1151 significantly lost peaks) compared to the limited gains of binding peaks (2 significantly gained peaks) in ZNF397-KO cells compared to control sgNT cells (Fig.4A, B, Fig.S6A, B, Table.S1), demonstrating ZNF397’s co-activator role in maintaining the AR cistrome. Consistent with this hypothesis, the most frequently lost binding motifs were those of the canonical luminal lineage driver transcription factors, including GR, AR, PGR, PR and FOXA1 binding motifs (Fig.S6C). To further confirm the crucial role of ZNF397 in maintaining the AR cistrome, we examined canonical AR target gene loci and revealed a profound loss of AR binding at those target genes (Fig. 4C, D: 14/20 AR Score Genes).

**Figure 4:**
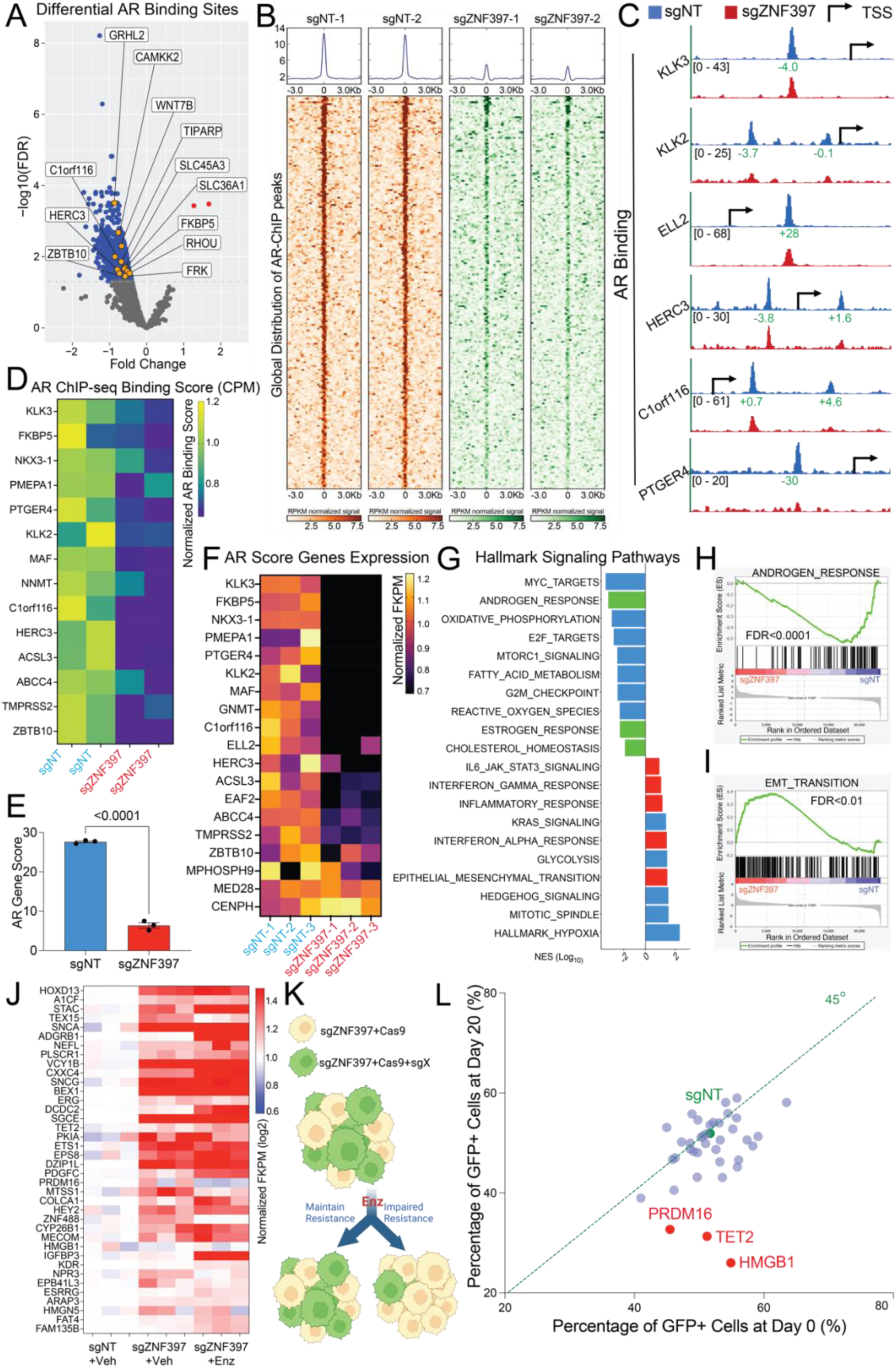
Integrated analysis of ChIP-Seq, RNA-Seq and CRISPR screening reveals key drivers of lineage plastic transcriptional programs. (A) Volcano plot represents the genomic loci with most significantly depleted or gained AR peaks, in ZNF397-KO cells compared to the control cells. Significantly changed gene loci were annotated as blue dots and identified AR Score Genes were annotated as yellow dots. Reads from two independently cultured cell samples were pooled for analysis. (B) Global distribution of AR binding peaks based on AR ChIP-seq results. Reads from two independently cultured cell samples were plotted. (C) Representative AR binding sites in the genomic loci of canonical AR gene loci in the LNCaP/AR cell transduced with Cas9 and annotated guide RNAs, based on AR ChIP-seq analysis. The binding peak distances (kilo base pair) to TSS (Transcriptional Start Site) are annotated in green and scales of peaks are annotated in black. (D) AR binding peak score (CPM, see Methods) in the genomic loci of AR Score genes (14 of 20) in the sgZNF397 cells compared to sgNT cells. Reads from two independent cell cultures/guide, matching input controls were used for analysis. (E)AR gene score based on the expression of canonical AR target genes (AR Score Gene) in ZNF397-KO and wildtype cells. mean ± s.e.m. is represented. p values were calculated using two-tailed t-test. (F) Heatmap represents the relative gene expression of AR Score genes in ZNF397-KO cells compared to wildtype cells, measured by RNA-Seq. (G) GSEA Pathways analysis shows cancer-related signaling pathways significantly altered in ZNF397-KO cells compared to wildtype cells, lineage specific pathways were highlighted with color: green-AR dependent and luminal lineage pathways, red-lineage plastic signaling pathways. (H) GSEA analysis of androgen response gene expression in ZNF397-KO cells compared to wildtype cells. (I) GSEA analysis of EMT-like lineage signature genes expression in ZNF397-KO cells compared to wildtype cells. (J) Unsupervised hierarchical clustering of normalized expression of differentially expressed genes whose expression was significantly changed in ZNF397-KO cells treated with vehicle or enzalutamide, comparing to sgNT+Veh group. 3 biological replicates in each group are shown. (K) Schematic representation of the functional CRISPR library screen in ZNF397-KO LNCaP/AR cells. sgZNF397 cells (GFP negative) were transduced with Cas9 and sgRNAs targeting individual candidate resistance driver genes (GFP positive). Then these sgZNF397+Cas9+sgX (GFP positive) cells were mixed with sgZNF397 (GFP negative) cells to achieve a cell mixture of 30-80% GFP positive cells. Schematic figure was created with BioRender.com. (L)Scatter plot summarizing the results of the screen. Each dot represents guide RNAs targeting a specific gene. The X-axis is the percentage of GFP cells at day 0 and the Y-axis is the percentage at day 20. The green dot identifies the sgNT control. Genes that scored positive in the screen are highlighted in red and labeled. Blue dot line represents the 45-degree line. For all panels unless otherwise noted, reads from 3 biological replicates were used for analysis. See also **Figure. S6**, **Table.S1-S4.**

RNA-Seq analysis revealed global changes in gene expression following the KO of ZNF397 (Fig. S6D, E, Table.S2), which were associated with changes in the AR cistrome. Consistent with the loss of AR binding to canonical targets, global transcriptomic analysis confirmed the substantial downregulation of canonical AR score genes following ZNF397-KO (Fig.4E, F). Notably, gene set enrichment analysis (GSEA) and pathway analysis revealed significant downregulation of the canonical AR signaling gene signature (Fig.4G, H, Table.S3). In contrast, this same analysis demonstrated enrichment in both the EMT-like lineage signature and signaling pathways known to promote lineage plasticity, including JAK-STAT, interferon response, and inflammatory response signaling (Fig.4G, I, Table.S3). The transcriptomic and cistromic findings confirmed the shift from an AR-dependent luminal state to an AR-independent lineage plastic state upon the loss of ZNF397.

Interestingly, upon loss of ZNF397 and abrogation of the AR-driven luminal lineage survival program, cell viability was maintained. Thus, we postulated that alternative transcriptional programs were activated and maintained the survival of PCa tumor cells. To this end, we examined both the transcriptomic and cistromic profiling results and identified the 38 most upregulated genes following ZNF397-KO (Fig.4J, Table.S4) as candidate alternative resistance drivers in the context of ZNF397 loss. To explore the functional role of how those candidates drive lineage plasticity and resistance, we utilized CRISPR deletion of each of these 38 genes to determine whether their deletion would impede the growth of enzalutamide resistant ZNF397-KO PCa cells (GFP+) (Table.S4). As assessed through a FACS-based competition assay (Fig. 4K; refer to Methods for details), the KO of three candidate resistance driver genes notably eliminated enzalutamide resistance in ZNF397-KO PCa cells (Fig. 4L, Table. S4). These genes included Ten-Eleven Translocation 2 (TET2), PR domain containing 16 (PRDM16), and high mobility group box 1 (HMGB1).

### TET2 is the crucial driver of lineage plasticity and AR-targeted therapy resistance

To validate the results of CRISPR library screening, we knocked out (KO) TET2, PRDM16 and HMGB1 in the ZNF397-KO cells and significantly impeded the resistant growth of those cells (Fig.5A, Fig. S6F), corroborating their crucial roles in mediating AR targeted therapy resistance. Among those three validated resistance drivers, the epigenetic modifier and 5hmC writer, TET2, plays a critical role in cell fate and lineage decision during both normal development and carcinogenesis^28–30,42,50^. Importantly, while TET2 is repressed by AR signaling in primary PCa, its expression and the TET2-dependent 5hmC are highly enriched in stem-like and neuronal lineages during normal prostate and PCa development ^37,51^. Therefore, the observed lineage transition from canonical luminal PCa to lineage plastic tumors in ZNF397-KO tumors raised the exciting possibility that TET2 is a master regulator conferring lineage plasticity and AR therapy resistance in PCa. Indeed, the expression of canonical TET2 target genes were substantially upregulated in ZNF397-KO cells compared to sgNT cells (Fig.5B), supporting an induction of TET2-driven transcriptional program ^52,53^. To assess whether TET2 is required for AR therapy resistance, we depleted TET2 from ZNF397-KO cells and revealed that TET2 depletion completely abolished the enzalutamide resistance conferred by ZNF397-KO, as shown in both FACS-based competition assay and cell viability assay (Fig. 5A, C). These results were corroborated by observations in a series of well-established patient-derived explant (PDE) models. In PDE samples treated with enzalutamide, significant downregulation of ZNF397 and AR target genes, along with upregulation of TET2 and its target ETS1, was observed compared to vehicle-treated samples (Fig. 5D, E).

**Figure 5:**
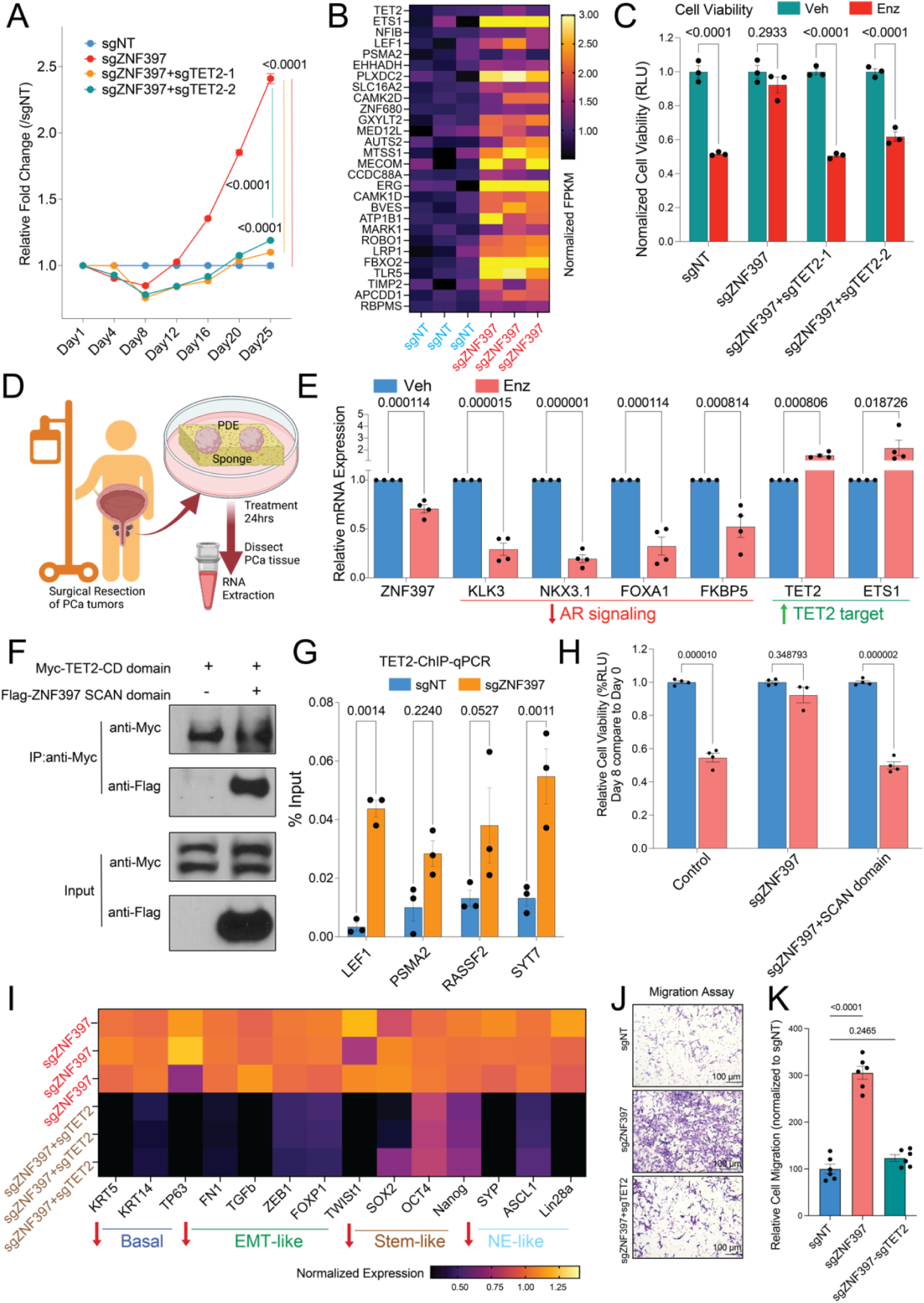
TET2 is the crucial driver of lineage plasticity and AR-targeted therapy resistance. (A) Relative cell number fold change of LNCaP/AR cells transduced with Cas9 and annotated CRISPR guide RNAs, measured by FACS-based competition assay. Enz denotes 10 µM enzalutamide treatment for 25 days. (B) Relative gene expression levels of canonical TET2 target genes in the LNCaP/AR cells transduced with Cas9 and annotated guide RNAs, measured by RNA-seq analysis. Data are normalized to the average expression in sgNT cells. (C) Bar plots represent the relative cell viability of LNCaP/AR cells transduced with Cas9 and annotated CRISPR guide RNAs, measured as values of relative luminescence unit (RLU) and normalized to vehicle treated conditions. Enz denotes 10 µM enzalutamide treatment for 7 days and Veh denotes DMSO. (D) Schematic figure represents the establishment of patient-derived explant models treated with vehicle or 10 µM enzalutamide for 24 hours. (E) Relative gene expression of ZNF397, AR pathway and TET2 target in independent patients derived explants (n=4) treated with vehicle (Veh) or enzalutamide (Enz) for 24 hours. p values were calculated using multiple t-tests with Benjamini correction. (F) Co-immunoprecipitation (Co-IP) of ZNF397 and the TET2 CD domain with a Flag or Myc tag in HEK293T cells. (G) TET2 ChIP-qPCR of canonical TET2 target genes’ genomic loci in LNCaP/AR cells transduced with annotated constructs. (H) Bar plots represent the relative cell viability of LNCaP/AR cells transduced with annotated CRISPR guide RNAs and rescue construct expressing ZNF397 SCAN domain, measured as values of relative luminescence unit (RLU) and normalized to vehicle treated conditions. Enz denotes 10 µM enzalutamide treatment for 7 days and Veh denotes DMSO. (I) Relative gene expression levels of canonical lineage specific marker genes in the LNCaP/AR cells transduced with Cas9 and annotated guide RNAs, measured by qPCR assay. Data are normalized to the average expression in sgNT cells. (J) Representative images of an LNCaP/AR cell transwell migration assay of six independent treated cell cultures. (K) Quantification of the number of migrated cells based on six representative images from three separate treated cell cultures for each cell line. For all panels unless otherwise noted, n = 3 independently treated cell cultures and mean ± s.e.m. is represented. p values were calculated using two-way ANOVA with Bonferroni multiple-comparison test. See also **Figure. S6**.

To dissect the molecular mechanism of how ZNF397-KO mediated TET2-driven resistance, we performed co-IP assays and observed robust interactions between the SCAN domain of ZNF397 and the cysteine-rich dioxygenase (CD) domain of TET2 (Fig.5F), which is consistent with a previous report that the ZNF397 SCAN domain blocks the DNA binding of the TET2 CD domain^54^. We then examined the TET2 binding on canonical TET2 target genes through TET2 ChIP-qPCR in ZNF397 deficient cells and observed significant upregulation of TET2 binding at those genomic loci (Fig.5G), suggesting that physiological level of ZNF397 functions as a barrier to TET2 DNA binding and function. This hypothesis is further supported by the observation that the resistance conferred by ZNF397-KO could be fully rescued by the reintroduction of the SCAN domain of ZNF397 (Fig.5H). We then assessed the expression of canonical lineage markers in the ZNF397 and TET2 double KO cells (sgZNF397+sgTET2). Strikingly, TET2 KO largely reversed the upregulation of basal, EMT-like, stem-like and NE-like marker genes (Fig.5I), which demonstrated its crucial role in maintaining lineage plastic transcriptional programs. The crucial role of TET2 in maintaining an EMT-like lineage program is further supported by the observation that the enhanced migratory abilities of ZNF397-KO cells were completely reversed upon TET2 KO (Fig.5J, K).

### Targeting TET2-driven epigenetic rewiring represents a novel therapeutic strategy to overcome resistance driven by lineage plasticity

Considering that TET2-dependent 5hmC modifications are highly correlated with prostate lineage decisions, we hypothesized that ZNF397-KO alleviates the blockade of TET2 at certain lineage plastic gene loci, thus activating these transcriptional programs through 5hmC modifications. To test this hypothesis, we evaluated TET2 binding to its target genes in ZNF397-KO cells, ZNF397/TET2-double-KO cells, and ZNF397/CXXC4-double-KO cells, as CXXC4 is responsible for recruiting TET2 to the sites of 5hmC modification ^55^. Remarkably, the deletion of CXXC4 entirely abolished the enhanced TET2 binding to its target genes (Fig.6A), suggesting that TET2-mediated resistance relies on 5hmC modification. To further corroborate this hypothesis, we assessed 5hmC modification on some known drivers of lineage plasticity, including SOX9, JAK1, and STAT1, and observed a significant induction of TET2-dependent 5hmC on these gene loci through 5hmC hydroxymethylated DNA Immunoprecipitation (hMeDIP) qPCR (Fig.6B). This effect is largely eliminated in the ZNF397/TET2-double-KO cells (Fig.6B).

**Figure 6.**
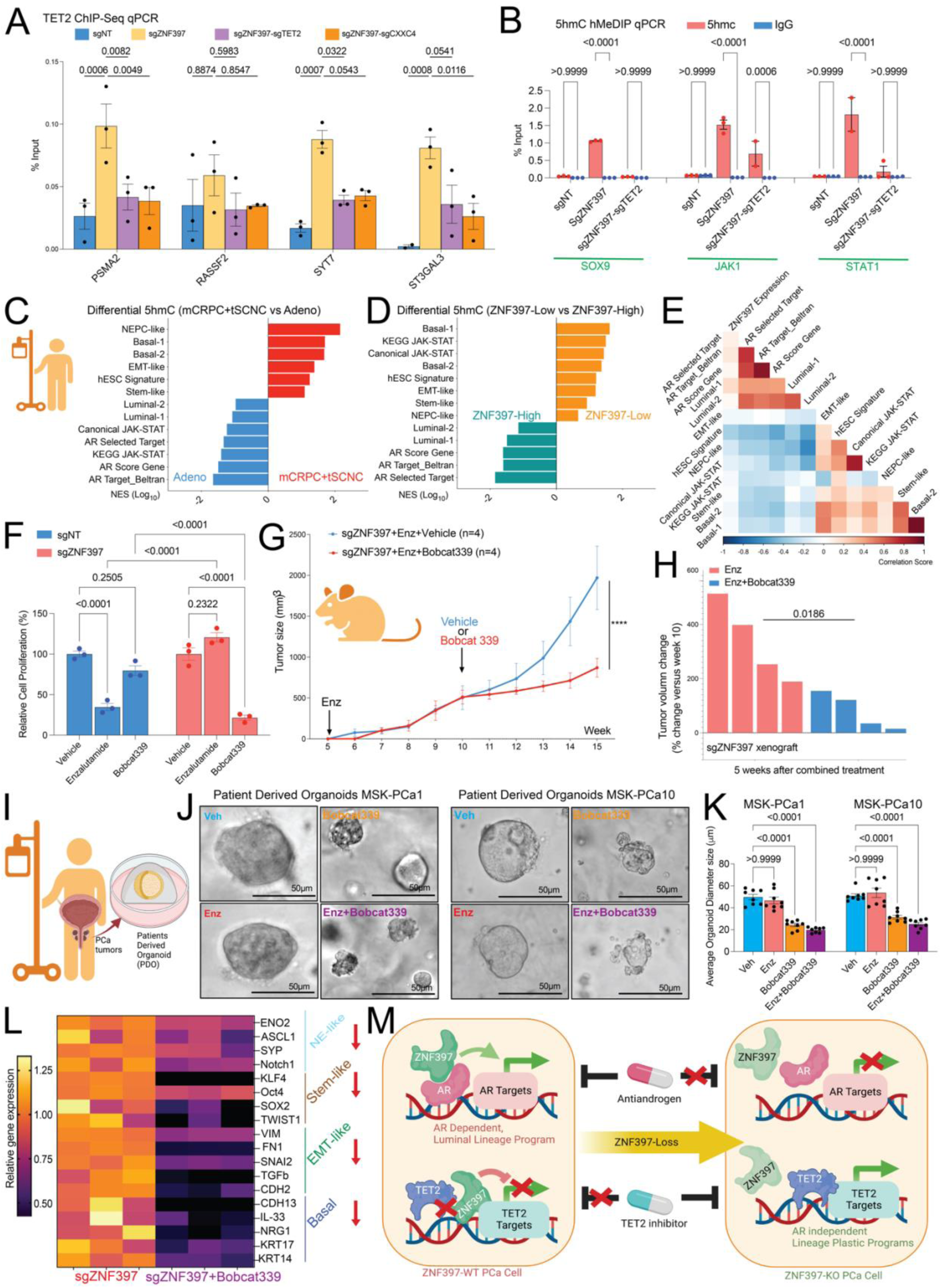
Targeting TET2-driven epigenetic rewiring represents a novel therapeutic strategy to overcome resistance driven by lineage plasticity. (A) TET2 ChIP-qPCR of canonical TET2 target genes’ genomic loci in LNCaP/AR cells transduced with annotated constructs. (B) 5hmC-hMeDIP-qPCR of lineage plasticity marker genes’ genomic loci in LNCaP/AR cells transduced with annotated constructs. (C) Bar plot presents the lineage gene signatures with most enriched or depleted 5hmC modification in the AR-independent mCRPC and t-SCNC patients compared to AR-dependent adenocarcinoma patients in a PCa patient cohort, based on the results of genomic 5hmC-Seq. (D) Bar plot presents the lineage gene signatures with most enriched or depleted 5hmC modification in the ZNF397-High patients (ZNF397 expression above median) compared to ZNF397-Low patients (ZNF397 expression below median) in a PCa patient cohort, based on the results of genomic 5hmC-Seq. (E) Heatmap represents the correlation between ZNF397 expression and the 5hmC modification in the genomic loci of those marker genes of lineage specific gene signatures, based on the results of genomic 5hmC-Seq. (F) Relative cell proliferation of LNCaP/AR cells transduced with annotated guide RNAs and treated with vehicle (DMSO), enzalutamide (10 μM for 7 days) or Bobcat339 (10 μM for 7 days). (G) Tumor growth curve of xenografted ZNF397-KO LNCaP/AR cells in castrated mice treated with enzalutamide. The mice were randomly separated into two groups when their tumors reached 500mm^3^ and treated with vehicle or Bobcat339. Enz denotes Enz treatment at 10 mg/kg from day 1 of grafting. Vehicle denotes denotes 0.5% CMC + 0.1% Tween 80, Bobcat 339 denotes Bobcat339 treatment at 10 mg/kg. n = number of independent xenografted tumors in each group. (H) Waterfall plot displaying changes in tumor size of xenografted ZNF397-KO LNCaP/AR cells after 5 weeks of combined treatments. Enz denotes Enz treatment at 10 mg/kg and Bobcat 339 denotes Bobcat339 treatment at 10 mg/kg. n = number of independent xenografted tumors in each group. (I) Schematic figure represents the establishment of patients derived organoid models MSK-PCa1 and MSK-PCa10 in 3D culturing system. (J) Bright field pictures represent the 3D-cultured patient derived organoid models, treated with vehicle (Veh), 5 µM enzalutamide (Enz), 10 µM Bobcat339 or enzalutamide+Bobcat339 for 7 days. (K) Bar plots represent the size of the 3D-cultured patient derived organoids, treated with vehicle (Veh), 5 µM enzalutamide (Enz), 10 µM Bobcat339 or enzalutamide+Bobcat339 for 7 days. (L) Relative gene expression levels of canonical lineage specific marker genes in ZNF397-KO and wildtype LNCaP/AR cells treated with vehicle (DMSO) or Bobcat339 (10 μM for 5 days), measured by qPCR assay. Data are normalized to the average expression in vehicle treated cells. (M) Schematic figure represents the innovative role of ZNF397 as a crucial co-activator of AR and co-suppressor of TET2. ZNF397-deficiency in prostate cancer promotes the transition of cancer cells from an AR-driven luminal lineage to a TET2-driven lineage plastic state expressing EMT-like, stem-like and NE-like lineages, which then fails to respond to AR-targeted therapy. See also **Figure. S6**, **Table.S5.**

A recent landscape study of prostate cancer 5hmC provided an opportunity to examine the crucial role of ZNF397 in global 5hmC modification and lineage plasticity ^37^. Using this study, we first examined the global distribution of 5hmC modifications in prostate cancers resistant to AR targeted therapy, including mCRPC and treatment-emergent Small Cell Neuroendocrine Cancer (t-SCNC), compared to AR-dependent prostate adenocarcinomas. As expected, 5hmC was substantially enriched at the signature genes driving lineage plastic transcriptional programs, including basal, EMT-like, stem-like, and NE-like lineages (Fig.6C, Table. S5) in the AR therapy-resistant tumors (mCRPC and t-SCNC). In contrast, 5hmC modifications were substantially depleted at the signature genes associated with an AR-dependent luminal lineage (Fig.6C), suggesting a 5hmC-related lineage switch abrogates AR dependency. We then divided the patients into two groups based on their ZNF397 expression and assessed the global distribution of 5hmC modifications. Strikingly, 5hmC modifications were substantially enriched at the lineage plastic gene loci but depleted at AR targets and luminal lineage gene loci in the ZNF397-low prostate cancers, compared to the ZNF397-high prostate cancers (Fig.6D), suggesting that ZNF397-deficiency is correlated with 5hmC rewiring and dynamically activated transcription of lineage plastic transcriptional programs. Finally, correlation analysis revealed that low ZNF397 expression is correlated with 5hmC enrichment at the lineage plastic signature genes, while high ZNF397 expression is correlated with 5hmC enrichment at the AR-driven luminal lineage signature genes (Fig.6E). Collectively, these clinical results validated the crucial role of ZNF397 in mediating 5hmC rewiring and lineage plasticity-driven therapy resistance.

Identification of TET2 as a critical mediator in the development of lineage plasticity-driven resistance raises hope that targeting TET2 and TET2-dependent 5hmC could overcome AR-targeted therapy resistance, providing a potential benefit to patients with advanced prostate cancer. Strikingly, the combination treatment of enzalutamide and the TET2 inhibitor, Bobcat339 ^56–63^, significantly impaired the growth of enzalutamide-resistant ZNF397-KO cells, as demonstrated in an *in vitro* cell viability assay (Fig.6F). Dose-response measurements (IC50) confirmed that ZNF397-KO cells were significantly more sensitive to Bobcat339 compared to wild-type cells (Fig.S6G). These *in vitro* results were further validated by *in vivo* xenograft experiments, as the combination treatment of Bobcat339 and enzalutamide stagnated the growth of Enz-resistant ZNF397-KO tumors (Fig.6G) and induced more potent tumor regressions than enzalutamide alone (Fig.6H). To further validate the clinical relevance of those results, we utilized two of the well-established patient-derived organoid (PDO) models MSK-PCa1 and MSK-PCa10 (Fig.6I), which have high expression of TET2 and treated those 3D-cultured PDOs with enzalutamide, Bobcat339 or the combination of those two agents. Strikingly, the enzalutamide resistant PDOs responded to Bobcat339 or the combination treatment of enzalutamide and Bobcat339 (Fig.6J, K), confirming the efficacy of targeting TET2 in overcoming enzalutamide resistance. Furthermore, Bobcat339 treatment profoundly abolished the induction of lineage plastic transcriptomic programs caused by ZNF397-KO, including basal, EMT-like, stem-like, and NE-like lineages (Fig. 6L).

Collectively, our results have revealed that ZNF397 is a crucial co-activator of AR and co-suppressor of TET2 in advanced prostate cancer. The deficiency (low expression) of ZNF397 in PCa is a molecular driver that promotes the transition of cancer cells from an AR-driven luminal lineage state to a TET2-driven lineage plastic state, which then fail to respond to AR-targeted therapy (Fig.6M). These findings uncover a cell-intrinsic molecular switch that determines tumor lineage plasticity and AR therapy resistance. Additionally, our findings highlighted TET2 as a promising therapeutic target for overcoming AR therapy resistance and validated the preclinical efficacy of a TET2 inhibitor in reversing lineage plasticity and AR therapy resistance, which may profoundly benefit patients with this devastating disease and other AR-dependent tumors.

## Discussion

Over the past decade, the development of a new generation of targeted therapies has revolutionized the management of many advanced cancers, including AR-targeted therapy for advanced PCa. Nevertheless, resistance to these targeted therapy agents in patients with metastatic PCa occurs rapidly and disease progression is often inevitable, drastically limiting the clinical outcomes of patients. Recently, emerging evidence has demonstrated that cellular plasticity, specifically lineage plasticity, functions as a predominant mechanism conferring targeted therapy resistance in various cancers, including prostate, breast, lung, and pancreatic cancers, as well as melanoma ^1^. Metastatic castration-resistant prostate cancer (mCRPC) stands as one of the most prominent illustrations of lineage plasticity, whereby luminal adenocarcinoma can transition to alternative lineage programs that are no longer sensitive to AR signaling blockage, such as neuroendocrine PCa (NEPC), double-negative PCa, or stem-like and multilineage phenotypes ^2^. Notably, significant dysregulation of the epigenetic regulation machinery often occurs during the acquisition of lineage plasticity, and several important epigenetic modifiers were implicated as key regulators of lineage plasticity in mCRPC, including EZH2, CHD1, and SOX2. However, the underlying mechanism of how cancer cells shift between different lineage programs through epigenetic rewiring remains largely unknown. Furthermore, there are no established therapeutic approaches available for patients with lineage plasticity-driven advanced cancers, highlighting the urgent and unmet clinical necessity for identifying druggable molecular drivers of lineage plasticity. Therefore, the identification of ZNF397 as a crucial molecular switch that governs the epigenetic rewiring and lineage plastic transition represents a major contribution of this work. Here, our results revealed that ZNF397 is a previously unrecognized key coactivator of the AR-driven luminal lineage survival program, while it is also a co-repressor for the TET2-driven non-luminal lineage survival program in advanced prostate and breast cancers. Since ZNF397 is frequently lost or downregulated in various cancers, the identification of this molecularly new subtype of aggressive cancers with ZNF397 deficiency and lineage plasticity not only clarifies our understanding of the origin and development of cancer therapy resistance but may also identify patients for early clinical intervention strategies to prevent or delay resistance.

In addition to the various genomic alterations already implicated in lineage plasticity, the dysregulation of epigenetic regulation machinery and epigenetic rewiring have been recognized as the predominant determinant of lineage plasticity and therapy resistance in various cancers ^6,8,22–26^. Among the many epigenetic modifications, cancer-associated DNA demethylation and 5hmC conversion, frequently mediated by TET2, exhibit a bifurcated role in the differentiation and progression of many advanced cancers. Interestingly, the function of TET2-mediated 5hmC transition in PCa is complicated and highly cell-type specific. On one hand, TET2 and TET2-dependent 5hmC oxidation is known to offset the transcriptional signaling driven by AR, which impairs the progression of primary PCa and is correlated with better clinical outcomes^34^.On the other hand, TET2-medaited 5hmC transition is highly enriched in the establishment of stem-like and neuronal lineages during development and carcinogenesis of various cancers ^43,44^ and, consequently, associated with poor clinical outcomes. Interestingly, TET2 was also implicated in the induction of cytokine signaling in T-cell-mediated immune responses ^50^, which was known to regulate PCa lineage plasticity ^17,18,64^. Contrary to the conventional view of its tumor suppressor role in primary PCa, our findings revealed that TET2-driven epigenetic reprogramming is a powerful mechanism that promotes the transition of mCRPC cancer cells from an AR-driven luminal lineage to a TET2-driven lineage plastic state expressing EMT-like, stem-like and NE-like lineages, which then fail to respond to AR-targeted therapy. These findings have revealed the unexpected oncogenic role of TET2 and have also mechanistically linked TET2-driven epigenetic rewiring to the acquisition of lineage plasticity and therapy resistance in prostate and other cancers.

Despite the clinical success of next-generation therapy agents targeting the AR signaling in PCa, resistance to these agents is inevitable and largely limits the clinical outcome of patients with advanced diseases ^65,66^. Recently, lineage plasticity has been authenticated as a widely utilized avenue to escape AR dependence and to promote AR-targeted therapy resistance in mCRPC ^2^. However, effective therapeutic approaches targeting lineage plasticity are not currently available, underscoring the necessity for identifying druggable targets which are crucial mediators of lineage plasticity. Here, the identification of TET2 as a key mediator of lineage plasticity and therapy resistance in mCRPC may represent an important advancement in the field of prostate cancer research. The ability to reverse lineage plasticity-driven resistance through genetic or pharmacological targeting of TET2 provides a potential avenue for developing effective therapeutic strategies for patients with advanced PCa. These findings may lead to the development of clinical trials to test the efficacy of TET2 inhibitors in treating mCRPC patients and overcoming resistance, thus contributing to improved clinical outcomes and overall survival. Collectively, this work highlights the importance of understanding the molecular mechanisms underlying lineage plasticity-driven therapy resistance and the potential of targeting these mechanisms for developing effective cancer therapies.

## Acknowledgments

We thank the SU2C, TCGA, cBioPortal.org, and Genomic Data Commons Data Portal (https://portal.gdc.cancer.gov/) for providing patients genomic and transcriptomic data. We thank Drs. Duojia Pan, Ralph Deberardinis, Michael Buszczak, Jun Wu and Kathryn O’Donnell for helpful discussion. This work was supported or partially supported by: National Cancer Institute/National Institutes of Health: 5R00CA218885 and 1R37CA258730 P.M., 1R01CA258584 T.W., T32C124334, 1F31CA261019-01A1 C.T.R.; Department of Defense: W81XWH-18-1-0411 and W81XWH21-1-0520 P.M., W81XWH2110418 XL.L.; Cancer Prevention Research Institute (CPRIT): RR170050, RP220473, P.M., RP230363 T.W.; Prostate Cancer Foundation:17YOUN12 P.M. 21YOUN10 M.S.; Welch Foundation: I-2005-20190330 P.M.; UTSW Deborah and W.A. Tex Moncrief, Jr. Scholar in Medical Research Award: P.M.; UTSW Harold C. Simmons Cancer Center Pilot Award: P.M.; UTSW CCSG Data Science Shared Resources: DSSR, T.W; Terry Fox New Frontiers Program Project Grant (PPG19-1090 to H.H.H.).

## Author contributions

Y.X. and P.M. conceived the project, with P.M. overseeing its progress, designing experiments, and interpreting data. P.M. authored the manuscript, while L.M. provided editorial assistance. Y.X., S.D., and N.J. cloned all plasmid constructs. Y.X. and N.J. conducted in vivo experiments, while Y.X., J.G., and S.D. performed *in vitro* cell-based assays. Y.X. established tumor-derived cell lines. Y.X. and N.J. executed immunohistochemistry (IHC). Y.X. and P.M. carried out and analyzed the CRISPR library screen. Y.X., J.G. and G.W. conducted western blots and qPCR. Y.W. and P.M. analyzed clinical data. M.H. and G.R. provided PDE samples. Y.C. provided PDO samples. Y.X. performed all PDE and PDO experiments. Z.W., S.B. and Y.W. conducted bioinformatic analysis of ChIP-Seq and RNA-Seq. T.W. and P.M. supervised bioinformatic analysis. M.S., H.H.H. and F.F. performed 5hmC analysis. Y.X., A.B.H., and C.A. executed breast cancer cell-related experiments. Y.X., X.Y., and Z.W. conducted co-immunoprecipitation (co-IP) experiments. H.Z. reviewed all statistical tests. P.M. is the corresponding author of this manuscript.

**Figure S1.**
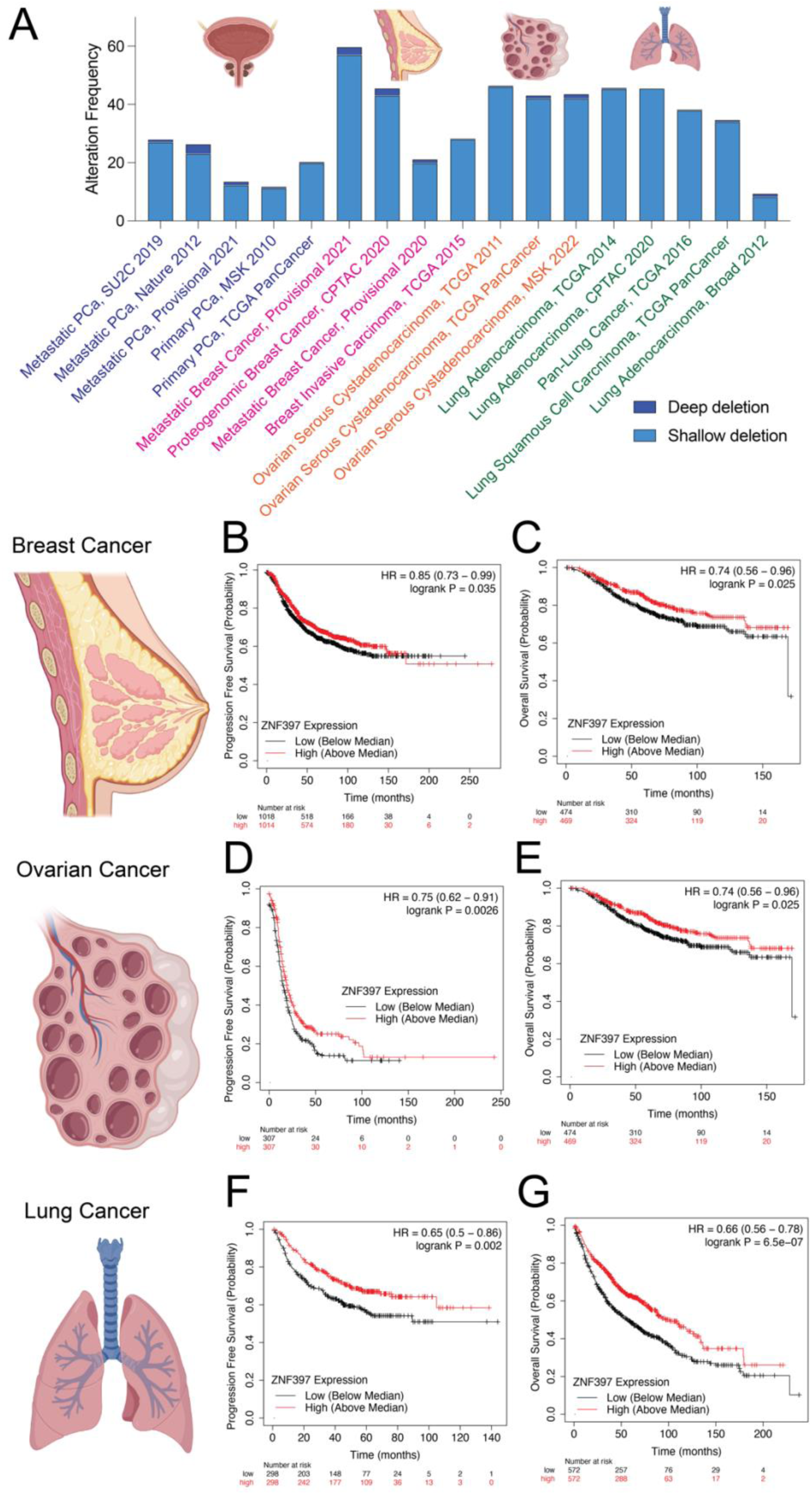
ZNF397-deficiency is correlated with poor clinical outcomes in various human cancers. (A) Stacked bar plot shows the percentage of cancer samples with genomic alterations within the ZNF397 locus in different patient cohorts with various cancer types, as created using cbioportal.org. (B-C) Kaplan-Meier plots represent the progression free survival (B) and overall survival (C) of breast cancer patients with high (above median) or low (below median) expression of ZNF397. (D-E) Kaplan-Meier plots represent the progression free survival (D) and overall survival (E) of ovarian cancer patients with high (above median) or low (below median) expression of ZNF397. (F-G) Kaplan-Meier plots represent the progression free survival (F) and overall survival (G) of lung cancer patients with high (above median) or low (below median) expression of ZNF397. For all panels, P values was calculated with Log-rank test and figure generated by Kaplan-Meier Plotter (see Methods). Schematic figures were generated with Biorender.com.

**Figure S2.**
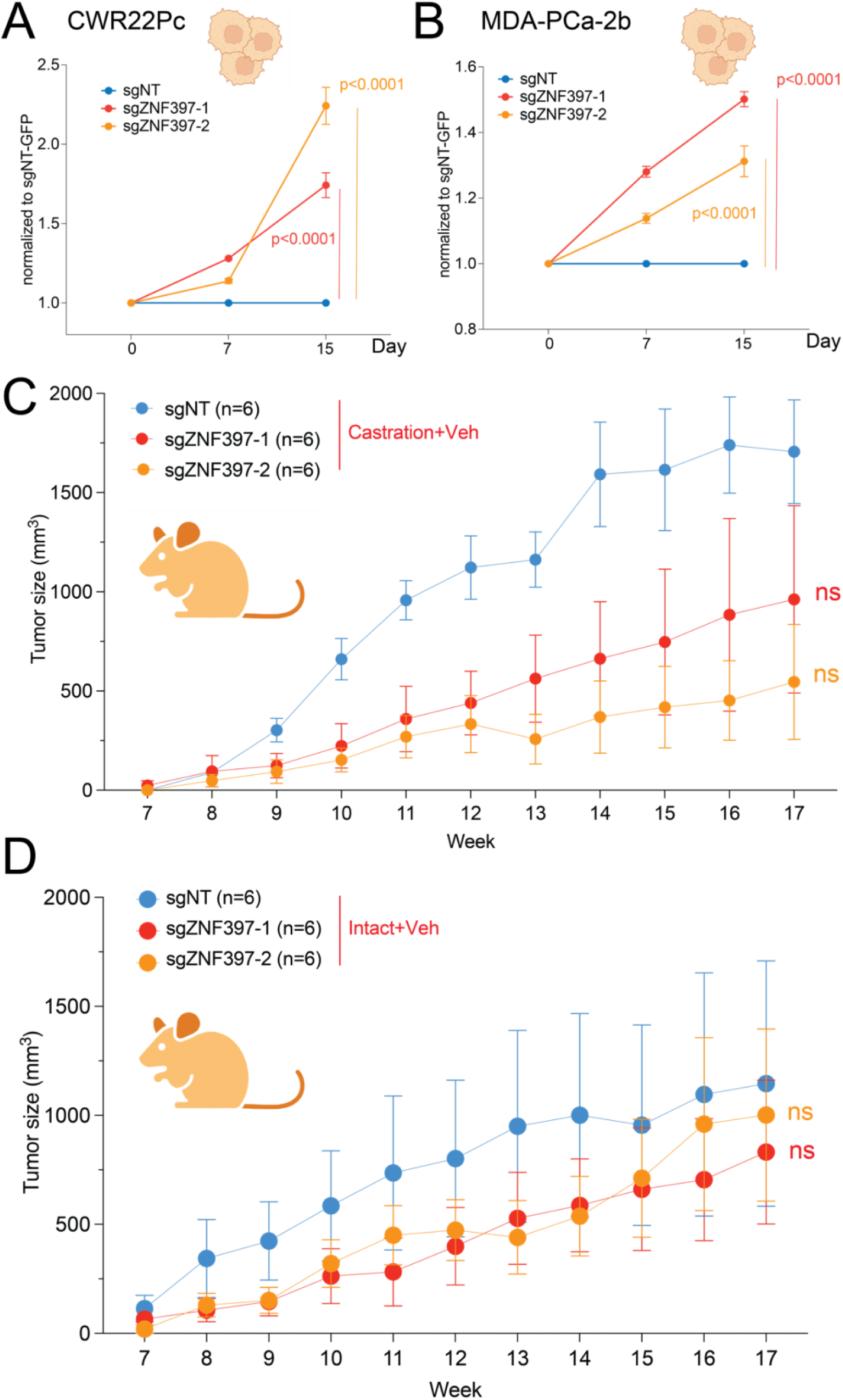
ZNF397-deficiency promotes antiandrogen resistance in other human PCa models. (A) Relative cell number fold change of CWR22Pc cells transduced with Cas9 and annotated CRISPR guide RNAs, measured by FACS-based competition assay. Enz denotes 1µM enzalutamide treatment for 15 days. (B) Relative cell number fold change of MDA-PCa-2b cells transduced with Cas9 and annotated CRISPR guide RNAs, measured by FACS-based competition assay. Enz denotes 10 µM enzalutamide treatment for 15 days. (C) Tumor growth curve of xenografted LNCaP/AR cells transduced with annotated guide RNAs in castrated mice treated with vehicle. Veh denotes denotes 0.5% CMC + 0.1% Tween 80. Number of tumors in each group were annotated. (D) Tumor growth curve of xenografted LNCaP/AR cells transduced with annotated guide RNAs in intact mice treated with vehicle. Veh denotes denotes 0.5% CMC + 0.1% Tween 80. Number of tumors in each group were annotated. Schematic figure was created with BioRender.com. For all panels unless otherwise noted, mean ± s.e.m. is represented and p values were calculated using two-way ANOVA with Bonferroni multiple-comparison test.

**Figure S3.**
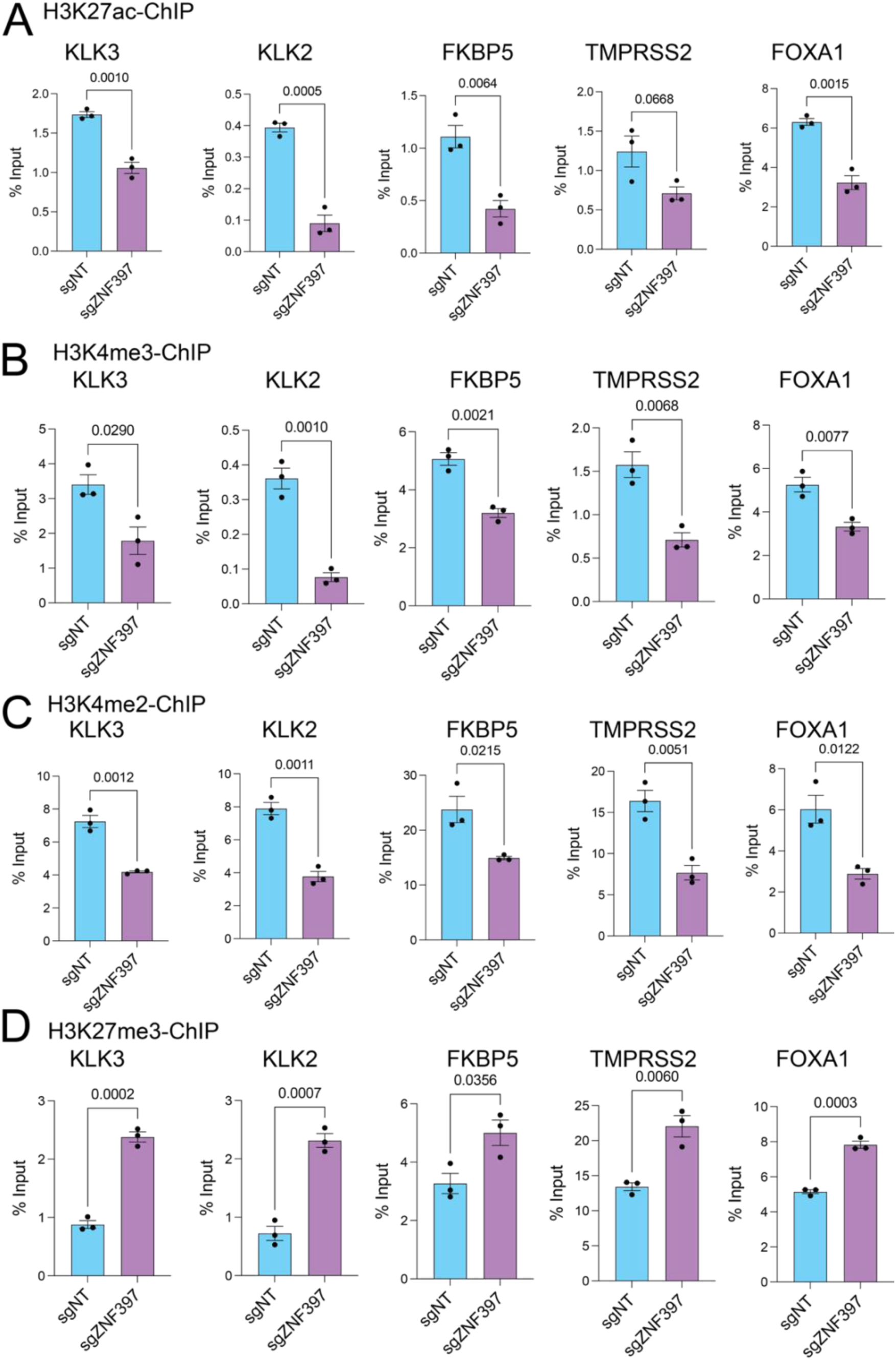
ZNF397 is required for the activation of canonical AR transcriptional program. (A) H3K27ac ChIP-qPCR of canonical AR target genes’ genomic locus in LNCaP/AR cells transduced with annotated constructs. (B) H3K4me3 ChIP-qPCR of canonical AR target genes’ genomic locus in LNCaP/AR cells transduced with annotated constructs. (C) H3K4me2 ChIP-qPCR of canonical AR target genes’ genomic locus in LNCaP/AR cells transduced with annotated constructs. (D) H3K27me3 ChIP-qPCR of canonical AR target genes’ genomic locus in LNCaP/AR cells transduced with annotated constructs. For all panels unless otherwise noted, n = 3 independently treated cell cultures and mean ± s.e.m. is represented. p values were calculated using two-tailed t-test.

**Figure S4.**
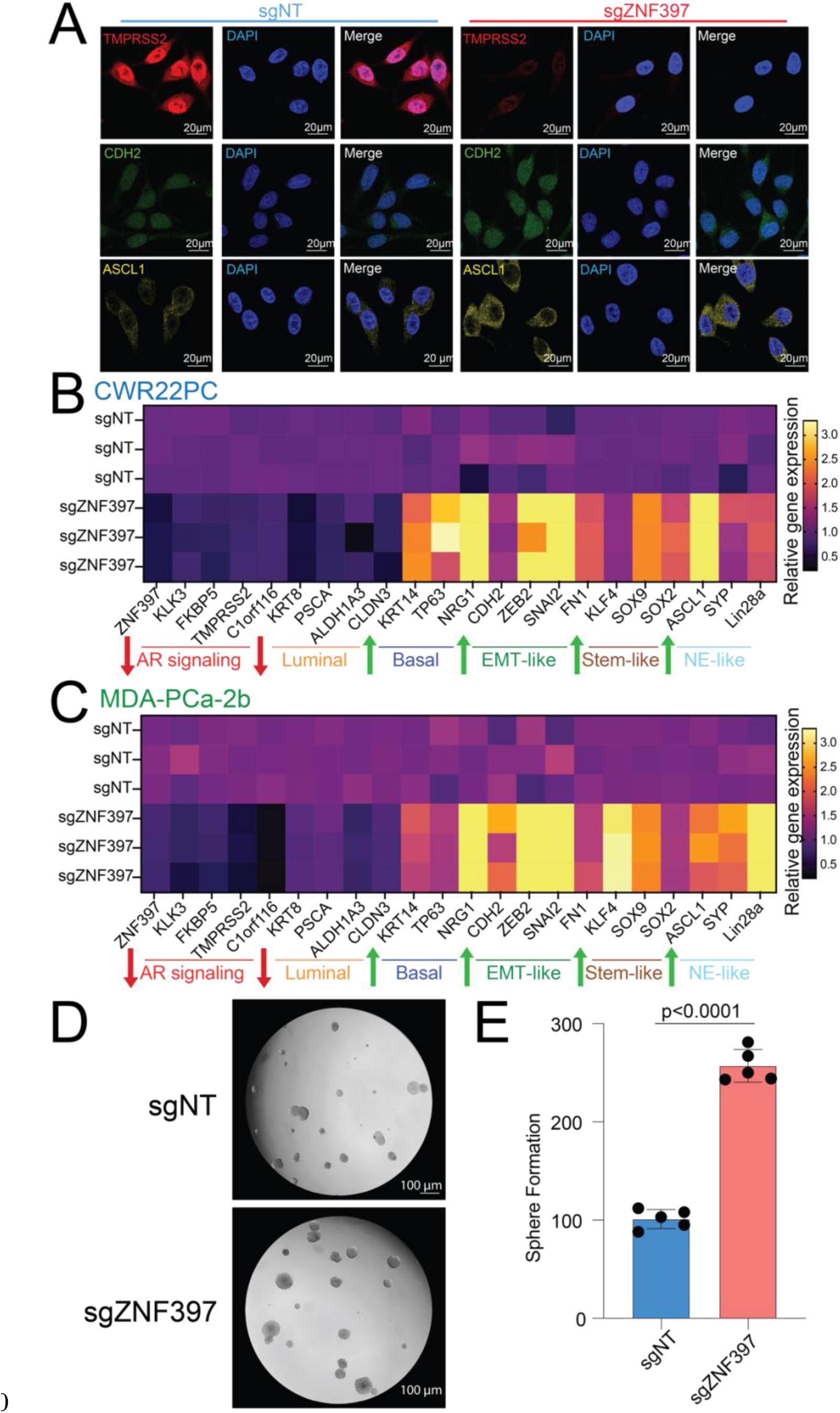
ZNF397-loss promotes lineage plasticity in two additional PCa models. (A) IF staining of LNCaP/AR cells transduced with Cas9 and annotated guide RNAs with annotated antibodies; representative pictures of n = 3 independently treated cell cultures were shown. (B) Relative gene expression levels of canonical lineage specific marker genes in the CWR22Pc cells transduced with Cas9 and annotated guide RNAs, measured by qPCR assay. Data are normalized to the average expression in sgNT cells. (C) Relative gene expression levels of canonical lineage specific marker genes in the MDA-PCa-2b cells transduced with Cas9 and annotated guide RNAs, measured by qPCR assay. Data are normalized to the average expression in sgNT cells. (D) Representative images of an LNCaP/AR cell prostasphere formation assay across five distinct treated cell cultures. (E) Quantitative analysis presenting the number of prostaspheres formed from five independently treated cell cultures for each cell line. Data are represented as mean ± s.e.m and P values were calculated by one-way ANOVA with a Bonferroni multiple-comparison test.

**Figure S5.**
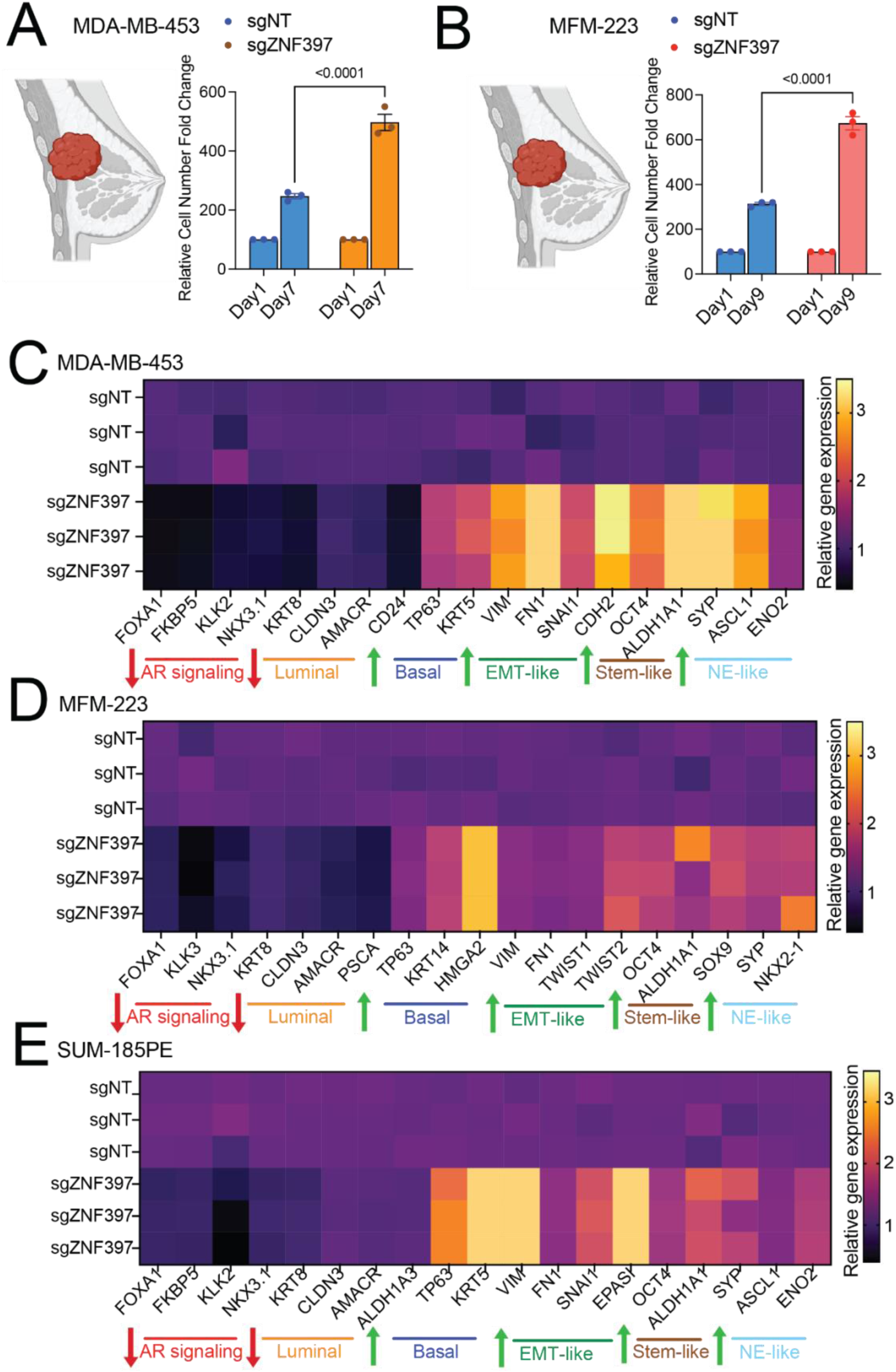
ZNF397-deficiency promotes tumorigenesis and lineage plasticity in breast cancer models. (A) Relative cell number fold change of MDA-MB-453 breast cancer cells transduced with Cas9 and annotated CRISPR guide RNAs, measured by FACS-based competition assay. (B) Relative cell number fold change of MFM-223 breast cancer cells transduced with Cas9 and annotated CRISPR guide RNAs, measured by FACS-based competition assay. (C) Relative gene expression levels of canonical lineage specific marker genes in the MDA-MB-453 breast cancer cells transduced with Cas9 and annotated guide RNAs, measured by qPCR assay. Data are normalized to the average expression in sgNT cells. (D) Relative gene expression levels of canonical lineage specific marker genes in the MFM-223 breast cancer cells transduced with Cas9 and annotated guide RNAs, measured by qPCR assay. Data are normalized to the average expression in sgNT cells. (E) Relative gene expression levels of canonical lineage specific marker genes in the SUM-185PE breast cancer cells transduced with Cas9 and annotated guide RNAs, measured by qPCR assay. Data are normalized to the average expression in sgNT cells. For all panels, data are represented as mean ± s.e.m and p values were calculated by two-way ANOVA with a Bonferroni multiple-comparison test.

**Figure S6.**
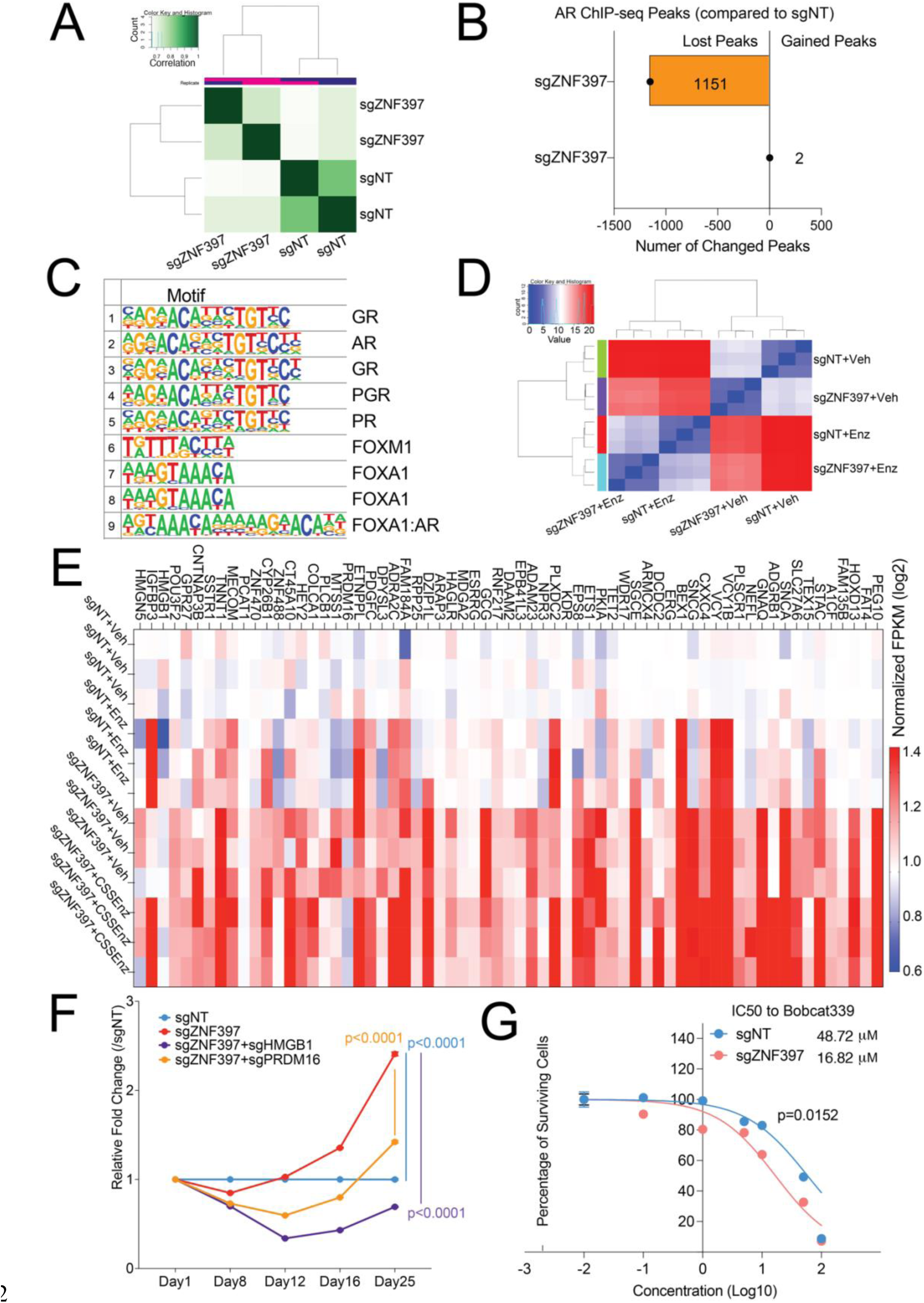
ZNF397-deficiency impairs the canonical AR cistrome and AR-driven transcriptional program. (A) Heatmap represents the correlation between the four samples in an AR ChIP-Seq analysis. (B) Number of significantly gained or lost AR peaks in ZNF397-KO cells compared to wildtype control cells. Reads from two independently cultured cell samples were pooled for analysis. (C) Motif enrichment analysis shows the most significantly lost motifs in the ZNF397-KO cells. The transcription factor binds to the corresponding motif are annotated on the right. (D) Heatmap represents the correlation between the transcriptional profiles of the ZNF397-KO and control cells, treated with vehicle or enzalutamide. (E) Heatmap represents the most significantly upregulated genes in ZNF397-KO cells compared to the control cells. (F) Relative cell number fold change of LNCaP/AR cells transduced with Cas9 and annotated CRISPR guide RNAs, measured by FACS-based competition assay. Enz denotes 10 µM enzalutamide treatment for 25 days. (G) Dose– response curve of ZNF397-KO and wildtype cells treated with TET2 inhibitor Bobcat339. For all panels unless otherwise noted, n = 3 independently treated cell cultures and mean ± s.e.m. is represented. p values were calculated using two-way ANOVA with Bonferroni multiple-comparison test.

## METHODS

### RESOURCE AVAILABILITY

#### Lead Contact

Further information and requests for resources and reagents should be directed to and will be fulfilled by the Lead Contact, Dr. Ping Mu.

#### Material Availability

All cell lines, plasmids, and other reagents generated in this study are available from the Lead Contact with a completed Materials Transfer Agreement if there is potential for commercial application.

#### Data and code availability

- All the described bulk RNA-seq data, ChIP-seq data have been deposited in the Gene Expression Omnibus (GEO, RRID: SCR_005012) under the accession numbers GSE230602, reviewer token is inetuecofpatnyt. Accession numbers are also listed in the key resources table. Original western blot images will be deposited at Mendeley and be publicly available as of the date of publication. Microscopy data reported in this paper will be shared by the lead contact upon request.
- All analysis in this manuscript was performed using open-source software. Bulk RNA-seq analysis was done using the QBRC Bulk RNA-seq pipeline and the codes were deposited in gitub. (https://github.com/QBRC/QBRC_BulkRnaSeqDE). GSEA statistical analysis was carried out with the R package ‘fgsea’ (v1.14.0). Customized codes used for ChIP-seq analysis were deposited in github (https://github.com/zwang0715/Xu_et_al_ChIPseq). Detailed analysis approaches were described in **QUANTIFICATION AND STATISTICAL ANALYSIS** session below.
- Any additional information required to reanalyze the data reported in this paper is available from the lead contact upon request.

## EXPERIMENTAL MODEL AND SUBJECT DETAILS

### Ethics statement

Animals were housed under humidity- and temperature-controlled conditions with a 12h light/12h dark cycle in the pathogen-free facilities at UT Southwestern Medical Center by the Animal Resource Center and were monitored closely to minimize discomfort, distress, pain, or injury throughout the course of the *in vivo* experiments. Animals were removed from the study and euthanized if any signs of pain and distress were detected or if the tumor volume reached 2,000 mm^3^. The *in vivo* xenograft experiments were performed as previously described ^8,64^, and described in detail in the following “METHODS DETAILS” section. All animals were divided into each experimental group at random, without prior designation. The xenograft tumor cells injection assay and follow-up tumor treatment were performed by one researcher, while tumor measurement and data analysis were performed by a different researcher to ensure the studies were run in a blinded manner. All procedures adhered to the guidelines provided by the Panel on Euthanasia of the American Veterinary Medical Association and the animal protocol was reviewed and approved by the Institutional Animal Care and Use Committee (IACUC) of UT Southwestern Medical Center (protocol#2019-102493). Male C.B-*lgh-1*^b^/lcr Tac-*Prkdc*^scid^ SCID mice were obtained from Taconic Biosciences.

### Cell lines

Human prostate cancer cell lines LNCaP/AR and CWR22Pc were cultured in RPMI-1640 medium (Life Technologies 11875135) containing 10% fetal bovine serum (FBS, Thermo Fisher Scientific 26400044), 1% GlutaMAX™ Supplement (Gibco 35050061), 1% penicillin-streptomycin (Sigma-Aldrich P0781), 1% HEPES (Thermo Fisher Scientific 15630080), and 1% Sodium Pyruvate (Fisher Scientific 11360070). Cells were passaged every 3-5 days at 1:3 and 1:2 as previously described ^8,64^. For hormone-starving, LNCaP/AR cells were cultured in RPMI-1640 medium containing 10% charcoal-stripped serum (CSS, Thermo Fisher Scientific 26400044), 1% GlutaMAX™ Supplement (Gibco 35050061), 1% penicillin-streptomycin (Sigma-Aldrich P0781), 1% HEPES (Thermo Fisher Scientific 15630080), and 1% Sodium Pyruvate (Fisher Scientific 11360070). Human prostate cancer cell line MDA-PCa-2b was purchased from ATCC (CRL-2422, RRID: CVCL_4748) and were cultured in Ham’s F 12K (Kaighn’s) Medium supplied with 20% FBS, 1% p/s, 25 ng/ml Cholera toxin (Sigma C8052), 10 ng/ml mouse epidermal growth factor (Fisher CB-40010), 0.005 mM phosphoethanolamine (Sigma P0503), 0.1 ng/ml hydrocortisone (Sigma H0135), 45 nM sodium selenite (Sigma#S9133), and 5 μg/ml human recombinant insulin (Fisher 12-585-014) and were passaged every 3-5 days at 1:4 as previously described ^8,64^.

HEK293T was purchased from ATCC (CRL-3216) and cultured in high glucose DMEM (Thermo Fisher Scientific 11965126) containing 10% fetal bovine serum (FBS), 1% penicillin-streptomycin (p/s), 1% HEPES, and 1% sodium and were passaged every 3-5 days at 1:3 and 1:2. Breast cancer cell lines MDA-MB-453, SUM185PE, MFM-223 were gifts from Carlos L. Arteaga lab at UT Southwestern Medical Center, all cells were cultured in high glucose DMEM containing 10% fetal bovine serum (FBS), 1% penicillin-streptomycin (p/s), 1% HEPES, and 1% sodium and were passaged every 3-5 days at 1:3 and 1:2. The identity of all cell lines were verified by human short tandem repeat (STR) profiling cell authentication at UT Southwestern genomic sequencing core every year and all cells were routinely tested for mycoplasma using MycoAlert^TM^ PLUS Mycoplasma Detection kit (Lonza, Cat#LT07-710).

### CRISPR-Cas9, shRNA and overexpression Plasmid

All the plasmids used in this study were lentiviral-based constructs and modified as described before ^8,64^. Specifically, CRISPR-Cas9 and guide RNAs were generated from all-in-one lentiCRISPRv2 (Addgene#52961, RRID: Addgene_52961), LentiCRISPRv2-GFP (Addgene#82416, RRID: Addgene_82416), LentiCRISPRv2-mCherry (Addgene#99154, RRID: Addgene_99154), pLKO5.sgRNA.EFS.RFP (Addgene#57823, RRID: Addgene_57823) and pLKO5.sgRNA.EFS.GFP (Addgene#57822, RRID: Addgene_57822) plasmids. A non-targeting gRNA was used as empty control. Benchling (RRID: SCR_013955, https://benchling.com) was used to design sgRNAs. shRNA sequences were cloned into shRNA constructs SGEP (pRRL-GFP-miRE-PGK-PuroR), SGEP (pRRL-GFP-miRE-PGK-PuroR), LT3CEPIR (pRRL-TRE3G-GFP-miRE-PGK-PuroR-IRES-rtTA3) and LT3GEPIR (pRRL-TRE3G-GFP-miRE-PGK-PuroR-IRES-rtTA3), which were originally obtained from the laboratory of J. Zuber at the Research Institute of Molecular Pathology. The sequence of sgRNAs/shRNA is listed in Table.S6 Key Resources Table. ZNF397-Full length and ZNF397-SCAN was amplified from HEK293T cDNA, TET2-CD domain was amplified from the TET2 full length construct^51^. The ZNF397-SCAN overexpression plasmid used for rescue assays was mutated at the PAM sequence with a synonymous mutation. All the sequences were cloned into pcDNA3.1 or pLenti-CMV-P2A-Blast, a gift from Rob lab at UT Southwestern Medical Center.

## METHOD DETAILS

### In vivo Xenograft tumor formation

All animal experiments were performed in compliance with the guidelines of the Animal Resource Center of UT Southwestern. The xenograft experiments were performed as previously described ^8,64^. Specifically, 2 × 10^6^ LNCaP/AR cells were suspended in the injection solution (50% Matrigel (BD Biosciences, 356237) and 50% growth medium). Cells mixture was then subcutaneously injected into the flanks of 7-week-old castrated male C.B-Igh-1b/Icr Tac-Prkdc^scid^ SCID mice on both sides. All 6-7 weeks old SCID mice were purchased from Taconic Biosciences and separated into each experimental group randomly, without prior designation. Mice were then treated daily with 10 mg/kg enzalutamide or vehicle (1% carboxymethyl cellulose, 0.1% Tween 80, 5% DMSO) by gavage feeding one day after the injection. Tumor size was measured weekly by using digital caliper when the tumor became measurable. For experiments depicted in Fig. 6G, 10 mg/kg enzalutamide (daily) and/or 10 mg/kg Bobcat339 (daily) were given after 5 weeks of Enz-only administration when tumors averaged around 500 mm^3^ in size. The tumor cell injection and follow-up tumor treatment were performed by one researcher, while tumor measurement and data analysis were performed by a different researcher to ensure the studies were run in a blinded manner. No statistical method was used to predetermine sample size, which was decided based on previously established protocol ^8,64^.

### Lentivirus production and CRISPR /shRNA/overexpression cell lines construction

Genomic modified cells were constructed by lentiviral infection as previously described with little modifications ^8,64^. 24 hours before transfection, 1.5 × 10^6^ HEK293T cells per well were seeded in 6-well plate. Lentiviral plasmids encoding Cas9/sgRNAs, shRNAs or ZNF397-SCAN domain overexpression, packaging plasmid psPAX2 (Addgene#12260, RRID: Addgene_12260) and the envelope plasmid pVSV-G (Addgene#138479, RRID: Addgene_138479) were mixed in Opti-MEM I Reduced Serum Medium (Thermo Fisher Scientific 31985062). Meanwhile, lipofectamine 2000 transfection reagent (Thermo Fisher Scientific 11668500) was diluted in Opti-MEM. The diluted plasmid and lipofectamine 2000 were then carefully mixed and incubated at room temperature for 20 mins. The mixture was added to HEK293T cells drop-wisely. The medium was changed 8-16 hrs after transfection. 24 hrs and 48 hrs after the medium change, the virus-containing supernatants were filtered with a 0.45 μM syringe filter and were saved for cell virus infection. 4 × 10^5^ LNCaP/AR cells per well were seeded in 6-well plate one day before. Medium was then changed twice with 50% fresh virus + 50% fresh medium + 5 μg/ml polybrene 24 hrs and 48 hrs after cell seeded. The lentiviral virus-containing medium was replaced with normal culture medium after 24 hrs. Cells were then selected with 2 μg/ml puromycin (Fisher Scientific ANT-PR-1) or 10 μg/ml blasticidin S (Thermo Fisher Scientific A1113903) for 5 d. All the shRNAs and sgRNAs sequence are listed in Table. S6 Key Resources Table.

### Luminescent Cell Viability Assay

Cell viability was measured by using CellTiter-Glo® luminescent cell viability assay kit (Promega, Cat#7570) according to the manufacturer’s instructions. LNCaP/AR (1000 cells per well) or CWR22Pc (20, 000 cells per well) or MDA-PCa-2b (4000 cells per well) cells were seeded in 96-well plate and treated with enzalutamide (10 μM for LNCaP/AR and MDA-PCa-2b, 1 μM for 22Pc) or vehicle for several days (7 days for LNCaP/AR, 5 days for CWR22Pc and MDA-PCa-2b). For 5α-Dihydrotestosterone (DHT) treated cell viability assay, cells were starved in RPMI-1640 containing 10% Charcoal Stripped FBS (CSS) medium for 3 days. 1000 Cells were seeded in 96-well plate and treated with increasing concentration of DHT or Vehicle (EtOH) for 7 days. 100 μl of CellTiter-Glo® Reagent was added to each well and mix contents for 12 minutes on an orbital shaker. The luminescence was recorded by the SpectraMax iD3 Multi-Mode Microplate Reader. To ensure the results were not biased by prior knowledge of the treatment groups, the cell viability was automatically measured by the SpectraMax iD3 plate. Treatments were conducted in biological triplicate and all experiments were repeated at least twice and achieved similar conclusions. No data points were excluded. Three biological triplicates were used and mean ± SEM were reported.

### Fluorescence-activated single cell sorting (FASC)-based competition assay

Cell resistance to enzalutamide was tested by FASC-based competition as previously described ^8,64^. CRISRP-Cas9-sgZNF397s-GFP or Dox-inducible shZNF397s LNCaP/AR cells were mixed in equal proportions with sgNT-RFP or shNT-RFP cells. The mixed cells were treated with either 10 μM enzalutamide or vehicle and collected by Attune NxT (version 4.2.1627.1) to measure the percentage of GFP/RFP cells on day 0, 3, 7, 12, 16, 16, 21 and 28. Relative cell number fold change was calculated as previously described ^8,64^. Dox-inducible cells were maintained in 250 ng/ml Dox throughout the whole experiment. To ensure the results were not biased by prior knowledge of the treatment groups, the cell number and percentage were automatically measured by the Attune NxT Acoustic Focusing Cytometer. All experiments were repeated at least twice and achieved similar conclusions. No data points were excluded. Three biological triplicates were used and mean ± SEM were reported.

### In vitro migration, invasion assay and wound healing assay

The migration and invasion were performed in 24-well Plate with 8.0 µm transparent PET membrane chambers (Corning, 353097). For invasion assay, chambers were coated with 30 μg extracellular matrix gel (Corning, 354234). 20, 000 LNCaP/AR cells with different genomic modification were resuspended in 300 µl Serum-Free RPMI and seeded into the chambers. 700 µl RPMI with 10% FBS serum was added into the 24-well plate. Cells were cultured for 48 hours at 37 °C with 5% CO_2_. Chambers were washed by PBS once, fixed in 4% PFA for 5 min and then methanal for 20 min. Cells were then stained in 1% crystal violet and sealed on slides for further quantification. For wound healing assay, 1.5 × 10^6^ LNCaP/AR cells with different genomic modification were seeded in 6-well plates and cultured for 24 hours at 37 °C with 5% CO_2_. 200 ul tips were used to generate the wound scratch. Cell scratches were then captured at 96 hours. All the pictures were captured by Leica DMi8 inverted microscope and quantified by using ImageJ (version 2.0.0, RRID: SCR_003070). For blinding purpose, the pictures were coded to blind researchers to treatment or genotype groups prior to data analysis to avoid bias.

### Prostasphere assay

The prostasphere assay was performed as previously described with small modifications (Lin *et al.*, 2019; Deng *et al.*, 2021). Two hundred cells per well were suspended in basic organoid medium supplemented with 20 ng/ml epidermal growth factor and 10 ng/ml basic fibroblast growth factor and were seeded into 96-well ultralow attachment plate (Corning 29443). For each condition, three wells were prepared for statistical analysis. Prostaspheres were imaged at one picture/well and quantified 12 days after seeding. All images were quantified by using ImageJ (version 2.0.0, RRID: SCR_003070).

### Patient derived explant (PDE) and patients derived organoid (PDO) experiments

PDE models were established in the Raj laboratory, where around 1 mm^3^ sample were embedded in sponge and cultured in RPMI-1640 media with 10% FBS, 1% penicillin-streptomycin solution, 0.01 mg/mL hydrocortisone, and 0.01 mg/mL insulin ^3^. PDEs were treated with 10µM enzalutamide or DMSO for 24 hours and then RNAs were collected. RT-qPCR were performed and analyzed using standard protocol describe in ***RNA Extraction and RT-qPCR***. PDOs were cultured in 3D Matrigel with typical human organoid medium as described before^67^. PDOs were passaged every 7 days at 1:3 by using trypsin or a sterile glass pipette. For treatment assay, PDOs were cultured in typical human organoid medium supplemented with 5 μM Enz and/or 10 μM Bobcat339. Images were captured by using Leica DMi8 microscope. All images were quantified by using ImageJ (version 2.0.0, RRID: SCR_003070). For blinding purpose, the pictures were coded to blind researchers to treatment or genotype groups prior to data analysis to avoid bias.

### RNA Extraction and RT-qPCR

Cells was lysed in Trizol (Ambion 15596018) and 1 μg extracted RNA was used for cDNA synthesis by using SuperScript^TM^ IV VILO^TM^ Master Mix (Thermo Fisher 11766500) per manufacturer’s instructions. RT-qPCR was performed by 2 × PowerUP^TM^ SYBR^TM^ Green Master Mix (Thermo Fisher, A25778). Briefly, 1-10 ng single-strand cDNA was used in one reaction and each reaction was conducted in triplicates. Data was analyzed by the delta delta Ct method (2-ΔΔCq) and all the target genes’ expression were normalized to reference gene. Heatmaps and Bars visualizing the gene differential expression were created by Graphpad Prism 9 Software (RRID: SCR_000306, https://www.graphpad.com/scientific-software/prism/) with expression fold change normalized to control cell lines (sgNT or shNT transduced LNCaP/AR). Primers used for qPCR are listed in Table.S6 Key Resources Table. All experiments were repeated at least twice and achieved similar conclusions. No data points were excluded. Three biological triplicates were used and mean ± SEM were reported.

### Protein extraction and Western Blot

Cell pellets were resuspended in Radioimmunoprecipitation Assay lysis buffer (RIPA buffer, 150 mM sodium chloride, 1.0% NP-40 or Triton X-100, 0.5% sodium deoxycholate, 0.1% SDS, 50 mM Tris, pH 8.0) supplied with proteinase and phosphatase inhibitors for 20 min on ice. The protein supernatants were collected by centrifuge (20,000 g for 10 minutes at 4℃) and quantified by using Pierce BCA Protein Assay Kit (Cat#23225). Protein lysates were denatured in 4 × Laemmli sample buffer (Biorad 1610747) with 2-mercaptoethanol at 95 ℃ for 5 minutes and 10 μg protein per sample was loaded on SDS-PAGE gel running in 1 × NuPAGE MES SDS buffer (Invitrogen, Cat#NP0002). Separated proteins were transferred to PVDF membrane in 1 × Blot Transfer buffer (Invitrogen, Cat#BT00061). Membranes were blocked in 5% non-fat milk for 30 minutes and then incubated with diluted primary antibody overnight at 4℃. Membranes were then washed in 1X TBST for 3 times and incubated with Horse Radish Peroxidase (HRP) conjugated secondary antibody for 1 hour at RT. The signal was developed into X-ray film in a dark room by using ECL (Thermofisher 32209) or SuperSignal West Pico PLUS (Thermofisher 34580). The following antibodies were used in western blot (also listed in Table.S6 Key Resources Table): Rabbit anti-ZNF397 antibody, Sigma-Aldrich#HPA026087, RRID: AB_2677389; Cyclophilin B (D1V5J) Rabbit mAb, CST#43603S, RRID: AB_2799247; Dykddddk Tag (D6W5B) Rabbit mAb, CST#14793, RRID: AB_2572291; Ty1 monoclonal antibody, Diagenode#C15200054; Myc Antibody (9E10), Santa Cruz Biotechnology#sc-40, RRID: AB_627268.

### H&E Staining, Immunohistochemistry (IHC) and Immunofluorescence (IF)

Mice xenograft tumors were harvested and were washed once with cold PBS, then were immediately fixed in 10% Neutral Buffered Formalin (StatLab Medical Products, 28600-1) at 4 °C overnight. Fixed specimens were processed and embedded in paraffin by the UT Southwestern Tissue Management Shared Resource core. Tumor blocks were sectioned at 5 μm on a standard rotary microtome (Leica, Germany). For H&E staining, slices were hydrated through xylene and a series of ethanol, and then were stained with Hematoxylin (Sigma-Aldrich, MHS1) and Eosin (Sigma-Aldrich#230251). For IHC staining, after deparaffinization and hydration, slides were heated in citrate sodium buffer (Sigma-Aldrich, W302600) and then were incubated in 3% hydrogen peroxide solution (Sigma-Aldrich, H1009) in methanol (Fisher Scientific, AC423950025). Slides were blocked with 3% BSA in PBST for 30 min at room temperature and then incubate with primary antibody (Rabbit Anti-ZNF397 antibody, Sigma-Aldrich, HPA026087, RRID: AB_2677389; Rabbit anti-Ki67, CST#9129, RRID: AB_2687446) overnight at 4°C. Slides were then incubated with Biotin-conjugated anti-rabbit IgG antibody (Jackson Immunoresearch, Cat#711-065-152, RRID: AB_2340593) and peroxidase Streptavidin (Fisher Scientific#NC9705430). The brown reaction signals were produced by using ImmPACT DAB Peroxidase (HRP) Substrate kit (Vector laboratories, SK-4105). Leica DMi8 microscope was used to capture the Images. For IF staining, LNCaP/AR cells were seeded onto round glass coverslips one day before. Cells were then fixed with 4% paraformaldehyde and permeabilized with 0.5% Triton X-100. Subsequently, the cells were blocked with 3% bovine serum albumin in PBS and incubated with the primary antibody overnight at 4°C. The cells were then incubated with Alexa Fluor-labeled secondary antibodies for 1 hour at room temperature, and nuclei were stained with DAPI. Images were captured using a Zeiss LSM 700 confocal laser-scanning microscope. The following antibodies were used for IF staining: Rabbit anti-ZNF397 antibody, Sigma-Aldrich#HPA026087, RRID: AB_2677389; Psa/klk3 (D6B1) XP® Rabbit mAb, CST#5365, RRID: AB_2797609; Ndrg1 (D8G9) XP® Rabbit mAb, CST#9485, RRID: AB_2721143; Nkx3.1 (D6D2Z) XP® Rabbit mAb, CST#92998, RRID: AB_2800197; TMPRSS2 Antibody (H-4), Santa Cruz Biotechnology#sc-515727, RRID: AB_2892118; Alexa Fluor 647 anti-cytokeratin 8(EP1628Y), Abcam#ab192468, RRID: AB_2890258; Alexa Fluor 647 anti-cytokeratin 14(EP1612Y), Abcam#ab192056; Vimentin (D21H3) XP® Rabbit mAb, CST#5741, RRID: AB_10695459; N-cadherin Antibody (13A9), Santa Cruz Biotechnology#sc-59987, RRID: AB_781744; Anti-MASH1/Achaete-scute homolog 1 antibody, Abcam#ab211327, RRID: AB_2924270; Alexa Fluor 647-conjugated AffiniPure goat anti-mouse IgG (H + L), Jackson ImmunoResearch#AB_2338902, RRID: AB_2338902; Alexa Fluor 647-conjugated AffiniPure goat anti-rabbit IgG (H + L), Jackson ImmunoResearch#AB_2338078, RRID: AB_2338078. All primary antibodies were diluted into 1:100 and all the secondary antibodies were diluted into 1:1,000. For blinding purpose, the pictures were coded to blind researchers to treatment or genotype groups prior to data analysis to avoid bias.

### Co-Immunoprecipitation (Co-IP)

5 × 10^5^ HEK293T cells were seeded into each well of a 6-well plate. 2 μg plasmids (pcDNA3.1-AR-Flag, pcDNA3.1-TY1-ZNF397-Full length, pcDNA3.1-ZNF397-SCAN-TY1, pcDNA3.1-ZNF397-SCAN-Flag or pcDNA3.1-Myc-TET2-CD domain) were mixed with 10 μl Lipofectamine® 2000 in 200 μl OPTI for 20 min at RT. The mixture was added evenly to the cells. 48 hours later, cells were lysed in 200 μl ice-cold mammalian IP lysis buffer (25 mM Tris-HCl, pH 7.4, 150 mM NaCl, 1 mM EDTA, 10% NP40) supplemented with 1 mM PMSF and 1 × Pierce protease and phosphatase inhibitor (Pierce^TM^, A32965) for 30 min at 4 °C with rotating. Protein supernatant was then collected by centrifugation at 20,000 g for 10 minutes and 30 μl sample was saved as Input. The rest supernatant was then incubated with 2 μl of pre-washed Anti-FLAG® M2 Magnetic Beads (Millipore#M8823) overnight at 4 °C with rotating. The following day, the Dynabeads magnetic beads in combination with any target were isolated by magnet (Invitrogen™ 12321D). Beads were washed three times with cold IP lysis buffer and boiled together with Input samples in 1 X SDS loading buffer + 1% β-ME at 95°C for 10 minutes after the final wash. Proteins were separated by western blot and signals were further captured by ECL reagent. Antibodies used in this assay including rabbit anti-Flag, rabbit anti-HA antibodies, mouse anti-TY1. All experiments were repeated at least twice and achieved similar conclusions.

### ChIP-qPCR

ChIP experiments were performed as previously described ^3^. LNCaP/AR cells with different genomic modifications were starved under regular RPMI-1640 containing 10% Charcoal Stripped FBS (CSS) medium for 3 days. 8 × 10^6^ cells were plated in 15 cm plate one day before and treated with 10 nM Dihydrotestosterone (DHT) or vehicle for 4 hours. Cells then were fixed with 1% PFA for 10 min and 0.125 M glycine for 5 min. Cell pellet were washed by cold PBS and resuspended in 300 ul ChIP Lysis Buffer with protease inhibitor. The DNA were sheared into 200-300 bp by Bioruptor® Pico, 1% sample was saved as input and the rest samples were incubated overnight with 5 μg antibody for each reaction in 4 degree. Dynabeads Protein G was added to each reaction the next day and incubate for 4 hours in 4 degree. Wash each sample sequentially with low salt wash buffer, high salt wash buffer, LiCl wash buffer and TE buffer. The DNA was eluted from the beads and reverse-crosslinked in 0.2M NaCl at 65 degree for 4 hours. The input DNA and ChIPed DNA were purified using MinElute PCR Purification Kit (Qiagen 28006) and the concentration was test by Qubit and ready for qPCR/ChIP sequencing. For ChIP-qPCR, DNA was amplified by 2 × PowerUP^TM^ SYBR^TM^ Green Master Mix (Thermo Fisher, A25778). Each reaction was conducted in triplicate and the enrichment percentage to corresponding input was calculated. The following antibodies were used in western blot (also listed in Table.S6 Key Resources Table): Anti-Androgen Receptor Antibody, Abcam#ab108341, RRID: AB_10865716),; Normal Rabbit IgG Polyclonal Antibody, Millipore Sigma#12-370, RRID: AB_145841 ; Anti-Histone H3 (acetyl K27) antibody, Abcam#ab4729, RRID: AB_2118291; Anti-Histone H3 (tri methyl K4) antibody, Abcam#ab8580, RRID: AB_306649; Anti-Histone H3 (di methyl K4) antibody, Abcam#ab7766, RRID: AB_2560996. Primers used for qPCR are listed in Table.S6 Key Resources Table. Three biological triplicates were used and mean ± SEM were reported, and experiments were repeated at least 2 times and achieved similar conclusions. No data points were excluded.

### hMeDIP-qPCR

hMEDIP experiments were performed by hMeDIP kit (Active Motif Cat#55010) according to the manufactory’s instructions. Briefly, 20 μg DNA was extracted from LNCaP/AR cells with different genomic modifications by the DNeasy Blood & Tissue Kit (Qiagen Cat#69504) and then sonicated into 200 to 600 base pairs by Bioruptor® Pico. 1 μg sonicated DNA immunoprecipitated with antibody against 5hmC, or negative control IgG and Protein G magnetic beads. After washing, the enriched DNA were eluted and then purified by MinElute PCR Purification Kit (Qiagen Cat#28006). DNA was amplified by 2 × PowerUP^TM^ SYBR^TM^ Green Master Mix (Thermo Fisher, A25778). Each reaction was conducted in triplicate and the enrichment percentage to corresponding input was calculated. Primers used for qPCR are listed in Table.S6 Key Resources Table. Three biological triplicates were used and mean ± SEM were reported, and experiments were repeated at least 2 times and achieved similar conclusions. No data points were excluded.

## QUANTIFICATION AND STATISTICAL ANALYSIS

### Statistical Methods

Statistical details of each experiment were showed in figure legends. Two-tailed t-test with Welch’s correction for unequal variances was used to compare two groups of independent datasets that fit normality and homoscedasticity. When normality and homoscedasticity were not satisfied, Mann Whitney U Test (nonparametric Wilcoxon Rank Sum Test) was used when comparing gene expressions between two patients’ groups. For comparisons involving more than two groups, one-way or two-way ANOVA and Kruskal-Wallis nonparametric ANOVA were used as appropriate. Mean ± s.e.m were reported, and p values were calculated and adjusted for multiple comparisons (Bonferroni or Benjamini correction) when applicable. For survival studies, the Kaplan-Meier method was used to estimate and plot the survival curve, and the log-rank test and Cox proportional hazard ratio analysis were used to evaluate differences in survival data among different groups. For all *in vitro* experiments, three biological replicates were performed except as noted in figure legends.

### Bulk RNA-seq

LNCaP/AR cells with different genomic modifications were treated with 10 μM enzalutamide or vehicle for 6 days before the total RNA was extracted using Trizol (Ambion, Cat 15596018) following manufacturer’s instruction. The extracted total RNAs were sent to UT Southwestern Next Generation Sequencing Core to perform bulk RNA-seq following the core’s sequencing instruction. Briefly, RNA concentration was determined by Qubit fluorometer and RNA integrity was determined by using Agilent 2100 Bioanalyzer, and samples with RNA integrity number (RIN) Score 8 or higher were used for library preparation. RNA libraries were prepared by using TruSeq Stranded Total RNA LT Sample Prep Kit from Illumina. Strand specific cDNA was synthesized from poly-A RNA and was then a-tailed and ligated with indexed adapters. Samples are amplified by PCR and purified with Ampure XP beads, following validation on the Agilent 2100 Bioanalyzer. Samples was quantified by Qubit before they were normalized and pooled. Then all samples were run as paired end read 150 nucleotides in length on the Illumina HiSeq 2500 using SBS v3 reagents. Alignment, quantification, and differential analysis were performed using the QBRC_BulkRnaSeqDE pipeline (https://github.com/QBRC/QBRC_BulkRnaSeqDE). Briefly, Alignment of reads to human reference genome (GRCh38) was done using STAR (v2.7.2b) (v2.7.2b)^68^. FeatureCounts (v1.6.4) was then used gene count quantification^69^. Differential expression analysis was performed using the R package DEseq2 (v1.26, RRID: SCR_015687)^70^. Cutoff values of absolute fold change greater than 2 and FDR<0.05 were used to select for differentially expressed genes between sample group comparisons. Gene Set Enrichment Analysis (GSEA) was carried out with the R package fgsea (v1.14.0) using the ’KEGG’ and ’WikiPathways’ libraries, and a custom library on transcriptional regulatory interactions (’TRRUST_Transcription_Factors_2019’) downloaded from the Enrichr database^71^.

### ChIP-seq

The ChIPed DNA and input DNA were prepared same as ChIP-qPCR. The DNA concentration was tested by Qubit 4 Fluorometer and the samples were assessed by Agilent 2100 Bioanalyzer. 10 ng DNA was used in library preparation, following NEBNext® Ultra™ II DNA Library Prep kit (NEB, Cat E7103). All samples were processed by End preparation, Adaptor ligation, U excision, PCR enrichment and clean up. The final products were qualified by Qubit 4 Fluorometer and assessed by Agilent 2100 Bioanalyzer. All samples were run on Illumina NextSeq 500 by 1 x 75bp (SR75) in CRI Sequencing Facility (Children’s Medical Center Research Institute at UT Southwestern). Raw sequencing reads from ChIP-Seq experiments were quality controlled using FastQC Tool (v0.11.9) and aligned to human reference genome assembly (GRCh38/hg38) using Bowtie 2 (v2.4.4, RRID: SCR_016368)^72^ with default parameters. Duplicated reads were removed using Picard MarkDuplicates (v2.26.11, https://broadinstitute.github.io/picard/, RRID: SCR_006525). ChIP-Seq peaks were called using MACS2 (v2.1.2)^73,74^. Input DNA was used as the control for peak calling. Genome browser tracks were generated using bamCoverage (v2.4.1) ^75^ with parameter “—normalizeUsingRPKM” and visualized using Integrative Genomics Viewer^76^. Motif analysis was performed using homer (v4.11)^77^. Differential analysis of peak enrichment and generation of volcano plot were performed using the ‘DiffBind’ R package v3.9.0 ^78^. ChIP-seq binding score is based on count-per-million (CPM) values of peak regions, which were calculated using FeatureCounts (v1.5.3) and normalized for heatmap plotting.

### 5hmC and correlation analysis

5hmC sequencing of mCRPC samples were performed and processed as previously reported ^37^. Gene body counts for TSS to TES in Gencode v.28 including intronic regions were extracted using featureCounts from the Rsubread R package and used for differential analysis. Differential analysis was performed for protein coding genes using the DESeq2 R package adjusted for metastatic tissue site and otherwise default parameters. GSEA was performed using the pre-ranked method implemented in the fgsea R package with genes ranked by the DESeq2 statistic for a curated set of well-established pathway signatures (Table.S5). Pathway level enrichment scores were calculated using the singscore R package with gene body counts first normalized to TPM and then ranked^79^. Correlation between sample level pathway scores, or ZNF397 expression, were calculated using spearman correlation and visualized using the corrplot R package.

### Analysis of Human Prostate Cancer Dataset

Processed 444 SU2C metastatic prostate cancer patient cohort data, including RNA-seq data and enzalutamide/abiraterone treatment data were downloaded from cBioPortal (RRID: SCR_014555, http://www.cbioportal.org/). The same subcohort of 51 patients^8^ with clinical response data were examined for progression free survival. The progression free survival figure was generated by Graphpad Prism 9 Software (RRID: SCR_000306, https://www.graphpad.com/scientific-software/prism/) using Mantel-Cox test. Cox hazards ratio analysis was performed using R-package ‘survminer’. Patients’ data analysis for breast, ovarian and lung cancers were queried using the Kaplan-Meier Plotter ^46–48^. All other patient genomic research cohorts were queried using cBioPortal (RRID: SCR_014555, http://www.cbioportal.org/).

**Table S1.** Significantly Altered AR ChIP-seq Peaks in ZNF397-KO Cells Compared to Control Cells

**Table S2.** Differentially Expressed Genes in ZNF397-KO Cells Compared to Control Cells

**Table S3.** Significantly Enriched or Depleted Signaling Pathways in GSEA Analysis

**Table S4.** Candidate Resistance Driver Genes in ZNF397-KO Cells

**Table S5.** Lineage-Specific and AR Target Gene Signatures

**Table S6.** Key Resources

